# Dissecting the stability determinants of a challenging de novo protein fold using massively parallel design and experimentation

**DOI:** 10.1101/2021.12.17.472837

**Authors:** Tae-Eun Kim, Kotaro Tsuboyama, Scott Houliston, Cydney M. Martell, Claire M. Phoumyvong, Alexander Lemak, Hugh K. Haddox, Cheryl H. Arrowsmith, Gabriel J. Rocklin

## Abstract

Designing entirely new protein structures remains challenging because we do not fully understand the biophysical determinants of folding stability. Yet some protein folds are easier to design than others. Previous work identified the 43-residue □ββ□ fold as especially challenging: the best designs had only a 2% success rate, compared to 39-87% success for other simple folds (1). This suggested the □ββ□ fold would be a useful model system for gaining a deeper understanding of folding stability determinants and for testing new protein design methods. Here, we designed over ten thousand new □ββ□ proteins and found over three thousand of them to fold into stable structures using a high-throughput protease-based assay. Nuclear magnetic resonance, hydrogen-deuterium exchange, circular dichroism, deep mutational scanning, and scrambled sequence control experiments indicated that our stable designs fold into their designed □ββ□ structures with exceptional stability for their small size. Our large dataset enabled us to quantify the influence of universal stability determinants including nonpolar burial, helix capping, and buried unsatisfied polar atoms, as well as stability determinants unique to the □ββ□ topology. Our work demonstrates how large-scale design and test cycles can solve challenging design problems while illuminating the biophysical determinants of folding.

**Significance:** Most computationally designed proteins fail to fold into their designed structures. This low success rate is a major obstacle to expanding the applications of protein design. In previous work, we discovered a small protein fold that was paradoxically challenging to design (only a 2% success rate) even though the fold itself is very simple. Here, we used a recently developed high-throughput approach to comprehensively examine the design rules for this simple fold. By designing over ten thousand proteins and experimentally measuring their folding stability, we discovered the key biophysical properties that determine the stability of these designs. Our results illustrate general lessons for protein design and also demonstrate how high-throughput stability studies can quantify the importance of different biophysical forces.

## Introduction

Improving our understanding of the determinants of protein stability (2–4) would accelerate biological, biomedical, and biotechnology research. In particular, computational models of protein stability are commonly used for a range of applications, including protein design (5–7), stabilizing naturally occurring proteins (8, 9), and predicting the effects of point mutants (10–12). However, all of these models have important limitations. For example, most computationally designed proteins made by experts fail to fold and function (13–15). Nonexperts avoid computational design techniques because they are not reliable. These challenges stem from our incomplete understanding of the biophysical determinants of folding stability and from the difficulty of encoding these determinants into computational models for practical applications.

Recently, we introduced a high-throughput approach to study protein folding stability that is particularly helpful for improving computational modeling and design. In our approach, we designed thousands of *de novo* proteins and measured their folding stabilities using a yeast display-based proteolysis assay coupled to next-generation sequencing (1). Several new studies have applied our methodology (16–19) as it has several advantages. First, measuring folding stability for thousands of proteins makes it possible to statistically quantify biophysical features that contribute to stability. Second, examining diverse sequences makes it easier to derive principles that are not specific to a particular protein context. Finally, assaying computationally designed proteins focuses the experimentation on the regions of sequence and structural space that are predicted to be low energy according to a particular computational model, which is especially useful for improving that model.

We previously used this approach to increase the success rate (i.e. fraction of designs that form stable, folded structures) of *de novo* miniprotein designs from 6% to 47% (1). Three different protein topologies could be designed very robustly (39-87% success), but a fourth topology (□ββ□, 43 residues) proved very challenging. Only 2% of □ββ□ designs folded into stable structures despite the simplicity of the structure and four repeated efforts to improve the design procedure (Fig. 1A). This suggested that our design procedure and stability model were missing something fundamental about the □ββ□ topology, and that this particular fold could be a useful model system for building a deeper understanding of folding stability. Here, we investigated this by asking two main questions. First, how can we improve our design procedure to obtain a large number of stable □ββ□ proteins for further analysis? Notably, there are no naturally occurring examples of the 43-residue □ββ□ fold for us to learn from, although this architecture is similar to the unusual 55-residue □ββ□ fold of the gpW protein from bacteriophage lambda (20). Second, how do the biophysical and topological features of different □ββ□ designs combine to determine each protein’s folding stability? We investigated these questions by designing and experimentally testing over ten thousand new □ββ□ miniproteins using our high-throughput approach. We also examined whether the structure prediction model AlphaFold 2 (21) could be applied to differentiate stable and unstable designs.

**Fig. 1.**
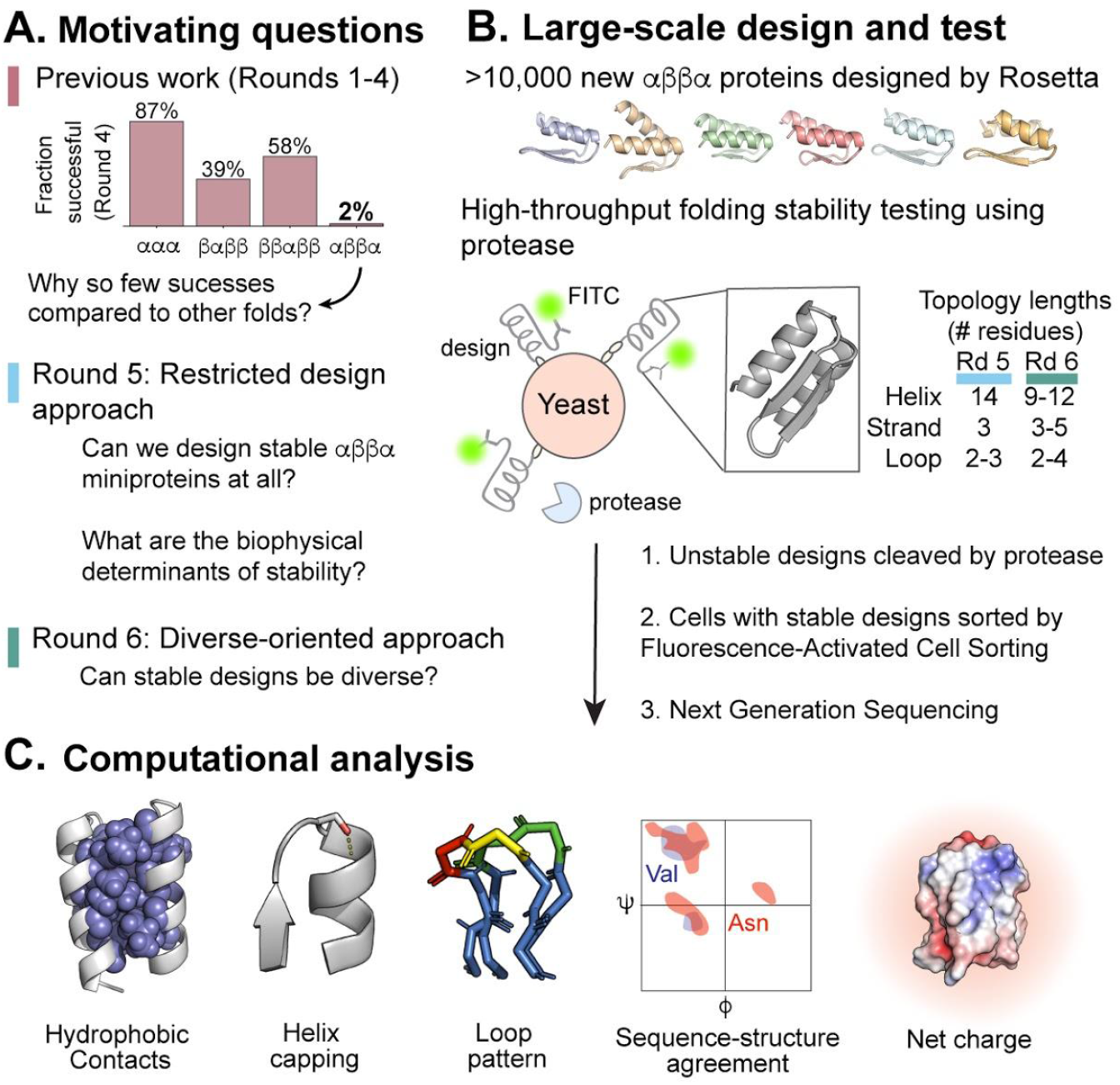
Design strategy for generating and testing αββα miniproteins. (A) Previously, we performed four iterative design-test-analysis cycles to generate stable αββα miniprotein designs, but only achieved a 2% success rate (1). (B) Here, we designed thousands of new αββα miniproteins using Rosetta (6,000 designs in Round 5 and 5,307 designs in Round 6) and experimentally tested them for their folding stability using a combined yeast display and protease sensitivity assay. (C) We then performed computational analysis to identify and understand the relative importance of key stability determinants (e.g. hydrophobic contacts, helix capping, loop patterning, local sequence-structure agreement, and net charge).

## Results

### Designing αββα miniproteins using a restricted design strategy

We first computationally designed thousands of new αββα miniproteins (“Round 5”) based on lessons learned from our previous four rounds of design (1). All designs were based on a single protein architecture (22) that previously led to the greatest number of stable designs (Fig. S1A). This architecture restricted our new αββα miniproteins to 14-residue α-helices, 3-residue β-strands, and a specific loop structure (Fig. 1B). In addition, we ensured our designs met strict criteria for buried nonpolar surface area, Rosetta energy, and predicted secondary structure (Fig. S1B). Finally, we required the middle loop to have a hydrophobic residue, required solvent-facing residues on the β-strands to be polar or charged, set a minimum threshold for the total number of hydrophobic residues, and eliminated Gly, Thr, and Val in helices (Fig. S1C-D) (see Methods). We hypothesized these restrictions would increase the success of our new designs because these constraints would enforce the overall αββα topology and build a larger hydrophobic core. However, this would reduce the potential sequence and structural diversity.

Based on this “restricted” design strategy, we generated 28,000 αββα miniproteins using an improved version of the Rosetta score function. This score function was previously parameterized to correlate with our earlier high-throughput data on miniprotein folding stability (23). In addition, we used an improved sequence sampling procedure that minimizes over-compaction and produces more native-like protein cores containing bulky residues (24). Our final set of 6,000 αββα designs were chosen by ranking the predicted stabilities of all 28,000 αββα designs using a linear regression model trained on previous large-scale αββα stability data (1). Because this regression model included a low number of stable designs (60/2830), we used this model for the practical task of selecting designs, but we did not expect reliable performance. After we ranked our designs, we eliminated designs that were more than 31/43 residues identical to a higher-ranking design. Within our final set of 6,000 designs, the median backbone RMSD between any two designs was 2 Å (Fig. S2) and the median sequence identity was 35% (Fig. S3). Each design based on this restricted strategy is named HEEH_TK_rd5_####, where HEEH indicates the pattern of α-helices (H) and β-strands (E), TK indicates the designer (author TEK), rd5 indicates these designs follow our four previous efforts (1), and #### is the design number.

### Biophysical characterization of αββα miniproteins using a restricted design strategy

We measured the folding stabilities of our newly designed αββα miniproteins using the high-throughput protease sensitivity assay introduced previously (Fig. 1B) (1). Briefly, all sequences were synthesized as DNA oligonucleotides in a pooled library. We then used *S. cerevisiae* to express and display our sequences on their cell surface, along with a c-terminal myc tag. Next, we subjected the yeast cells to varying concentrations of trypsin and chymotrypsin (tested separately) (Fig. S5A-B) and fluorescently labeled the cells displaying protease-resistant sequences. Finally, we sorted the fluorescently labeled cells by flow cytometry and identified the protease-resistant sequences by deep sequencing (Fig. 1B). Out of the 6,000 designs, only 5,662 designs had sufficient sequencing counts to precisely determine their protease sensitivity, and we used this set of 5,662 designs for our analysis. As previously, we assigned each design a “stability score”, defined as the difference between that sequence’s observed protease sensitivity and the predicted sensitivity of that sequence in its unfolded state. Each one-unit increase in stability score indicates a 10-fold higher amount of protease required to cleave that sequence under assay conditions, compared with the predicted protease concentration required to cleave that sequence in its unfolded state (1). To conservatively identify stable designs, each design’s overall stability score is the minimum of the stability scores observed separately with trypsin and chymotrypsin. We previously observed that sequences of scrambled amino acids (not designed sequences) rarely have stability scores above 1, and so we classify designs as stable when their stability score exceeds 1.

Our set of 5,662 designs had an average stability score of 0.81, and we classified 38% of these designs as stable (stability score > 1, Fig. 2A). The stable set had a median pairwise sequence identity of 37% (Fig. S3). This greatly exceeded our previous success rate of 2% (Fig. 1A) (1). We also included control sequences in our library whose residue compositions matched our αββα designs, but with the ordering of the residues scrambled in a specific manner: polar residues remained polar, nonpolar residues remained nonpolar, and proline and glycine residues remained in their identical positions. In contrast to our designs, almost all scrambled sequences had stability scores < 1 with an average stability score of −0.86 (Fig. 2A). This suggests that the protease resistance observed for a subset of designs can be attributed to the folding stability of their designed structures, rather than generic properties of their sequences such as residue composition or patterning. In addition, stability scores measured using trypsin and chymotrypsin were correlated with each other despite the differing specificities of the proteases (Fig. S2A-B). This further indicates that our measured stability scores reflect folding stability rather than protease-specific factors.

**Fig. 2.**
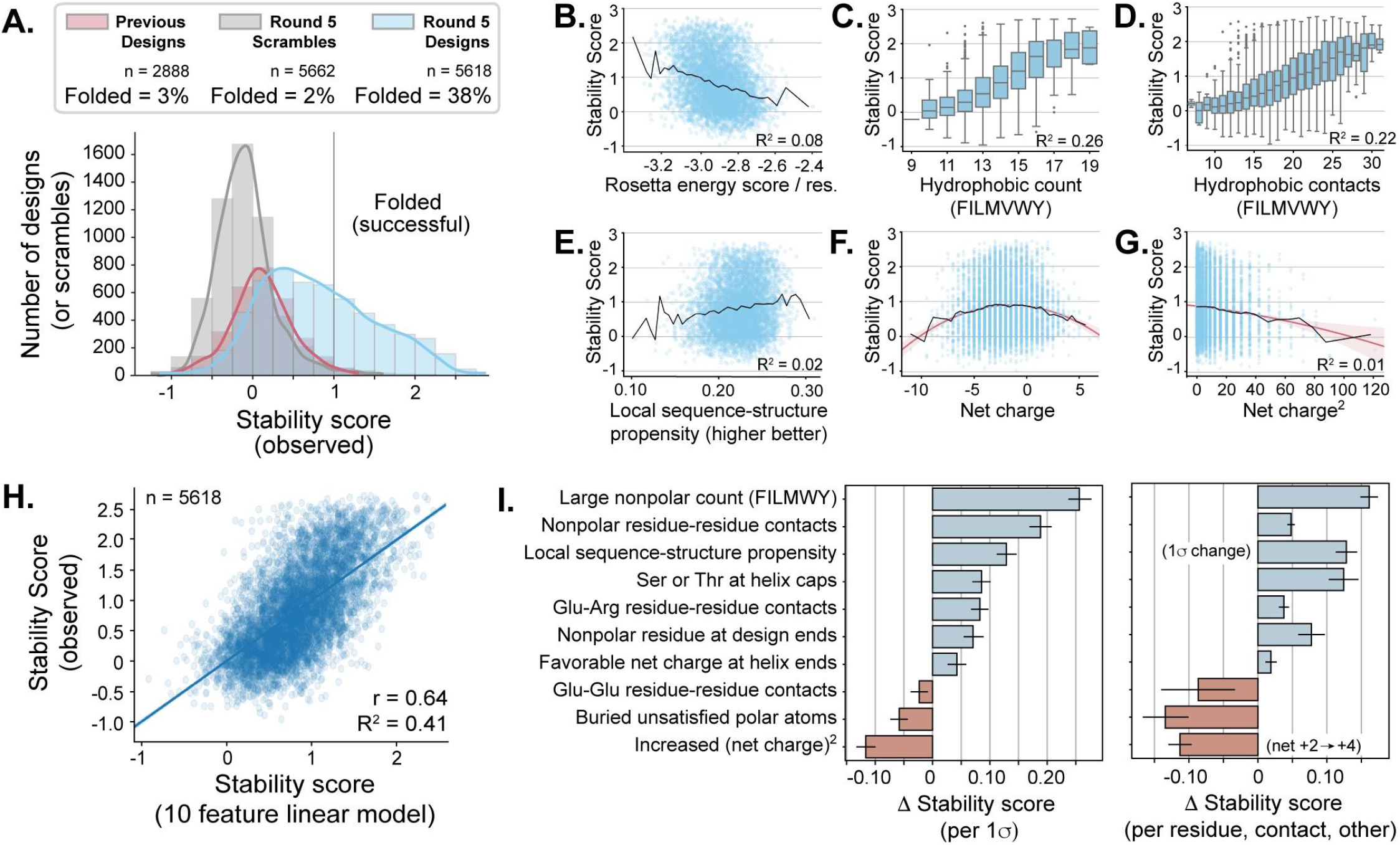
Experimental testing and analysis of αββα stability determinants from a restricted design strategy. (A) The stability score distributions of designed αββα miniproteins (blue), scrambled sequences (gray) and previously published αββα miniproteins (red) (1); the vertical line at stability score = 1 denotes the threshold above which we consider a design to be stable. (B-G) The relation between Individual protein features and stability score. For Rosetta energy, lower values indicate favorable energies, and for local sequence-structure propensity, higher values indicate favorable propensity. Black lines show moving averages; red lines show fits to quadratic (F) and linear (G) models. (H) A ten-feature linear regression model was built using normalized data, and the experimental stability scores are compared to the model’s predicted stability scores. (I) The magnitudes of the coefficients from the model based on their importance in the dataset (left) and their biophysical strength (right). Error bars indicate 95% confidence intervals from bootstrapping.

We next sought to verify that stable αββα miniproteins folded as designed using several orthogonal approaches. First, we selected six stable αββα designs with varying hydrophobicity values (25) and individually purified them from *E. coli* (Fig. 3A; Table S1) for circular dichroism (CD) and thermal denaturation. Protein purification by size-exclusion chromatography revealed that three of the six miniproteins (HEEH_TK_rd5_0420, HEEH_TK_rd5_0614, and HEEH_TK_rd5_0958) predominantly eluted at the expected molecular weight of a monomer, whereas the other three showed both monomeric and dimeric peaks (Fig. 3B).

**Fig. 3.**
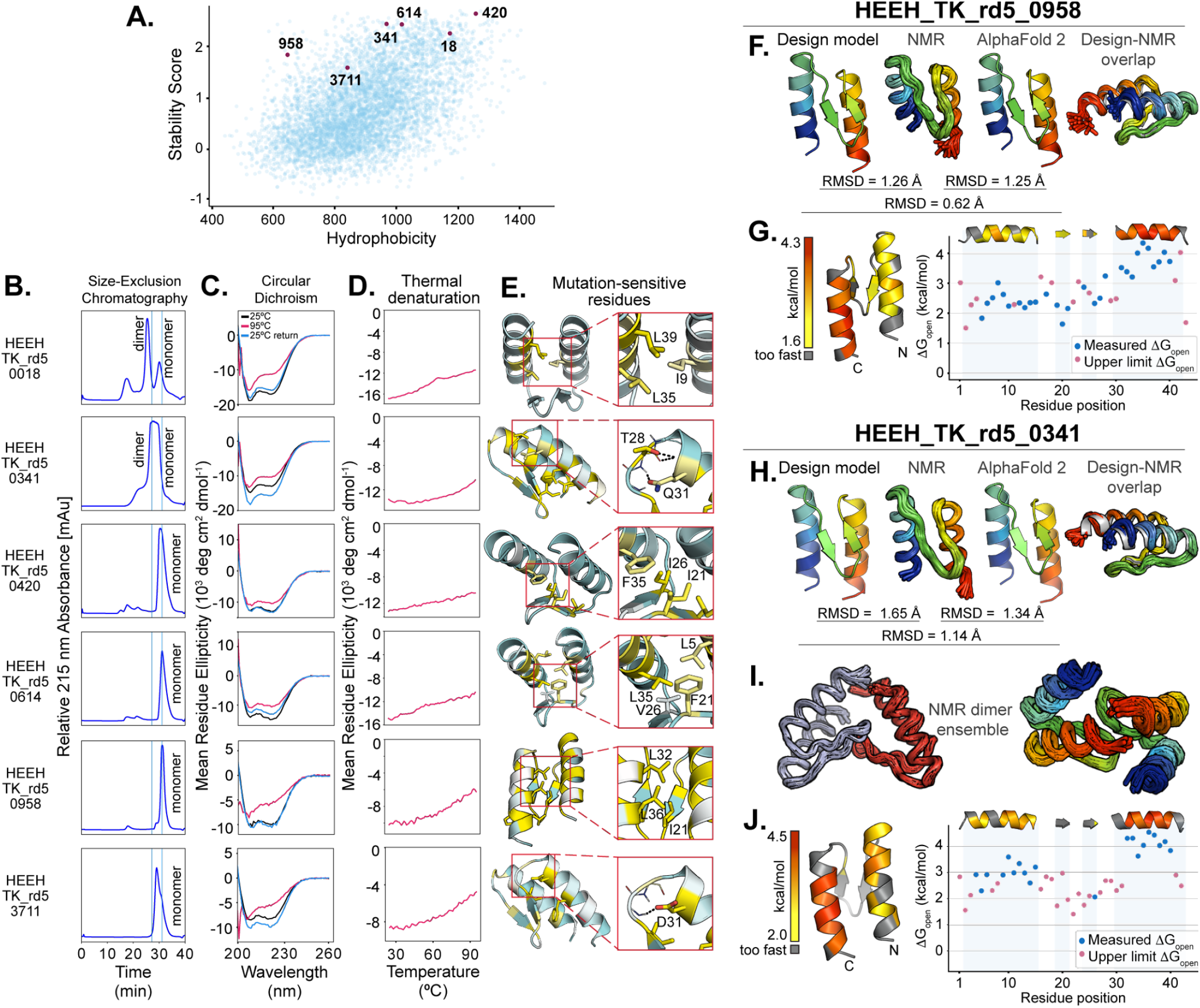
Biophysical characterization of αββα miniproteins made using a restricted design strategy. (A) The stability scores of all αββα miniproteins made using a restricted design strategy are plotted by their hydrophobicity values (27). We selected six miniproteins (red dots) with varying hydrophobicity and (B) purified them by size-exclusion chromatography; vertical lines indicate expected dimeric and monomeric forms of the miniprotein based on a calibration curve (see Methods). (C) Far-ultraviolet circular dichroism spectra are shown at 25°C (black), 95°C (red), and 25°C after melting (blue). (D) Thermal denaturation was measured at 220 nm at every 1°C from 25°C to 95°C. (E) Design models highlight positions that are most tolerant (teal) or least tolerant (yellow) to mutations. Key residues that stabilize the protein are shown in stick representation. Each miniprotein’s color scale is different to highlight the relative stabilizing or destabilizing effects within each protein; see Fig. S5 for complete data. (F) Comparison of HEEH_TK_rd5_0958 Rosetta design model, NMR ensemble, and AlphaFold 2-predicted structures; overlay of the Rosetta design model (gray) and NMR ensemble (rainbow). (G) Opening energies determined by hydrogen-deuterium exchange for HEEH_TK_rd5_0958. Observed measurements are colored red-yellow on a cartoon model and plotted in blue. For residues that exchanged too quickly to measure, the upper limit of ΔG_open_ is plotted in red. (H) Comparison of HEEH_TK_rd5_0341 Rosetta design model, NMR ensemble, and AlphaFold 2-predicted structures; overlay of the Rosetta design model (gray) and NMR ensemble (rainbow). (I) NMR dimer structure shown in two different perspectives. (J) Opening energies determined by hydrogen-deuterium exchange for HEEH_TK_rd5_0341. Observed measurements are colored red-yellow on a cartoon model and plotted in blue. For residues that exchanged too quickly to measure, the upper limit of ΔG_open_ is plotted in red.

CD spectra exhibited helical secondary structure and reversible folding after heating to 95°C (Fig. 3C), but the initial 25°C and cooled 25°C return measurements for HEEH_TK_rd5_0341 and HEEH_TK_rd5_3711 were not superimposable, possibly due to aggregation that altered the signal intensity. None of the designs showed a clear melting transition, although designs HEEH_TK_rd5_0958 and HEEH_TK_rd5_3711 lost much of their helical character at 95°C. In contrast, design HEEH_TK_rd5_0420 was minimally perturbed during melting (Fig. 3D), indicating extreme thermostability.

Next, to spot-check the accuracy of our designed structures, we solved the structures of HEEH_TK_rd5_0958 and HEEH_TK_rd5_0341 by nuclear magnetic resonance (NMR). For HEEH_TK_rd5_0958, the average backbone root-mean-squared deviation (RMSD) of the design model compared to all 20 structures in the NMR ensemble was 1.26 Å (Fig. 3F). For HEEH_TK_rd5_0341, both monomers in the dimeric structure were also very close to the designed monomeric model: the average backbone RMSD of the design model compared to all 40 structures in the NMR ensemble was 1.65 Å (Fig. 3H). The structure was symmetrical so only one HSQC peak was visible for each residue, although ^15^N NMR relaxation measurements were consistent with the dimeric state. The two monomers come together near the β-hairpin and designed N- an C-termini, burying hydrophobic residues in that region (Fig. 3I).

To analyze the structural differences between the design models and NMR structures, we quantified the number of contacts a residue in the design model gained or lost in the NMR model (Fig. S6). Most of the residues gained or lost zero or one contact, indicating the close structural similarity between the design model and NMR ensemble. For HEEH_TK_rd5_0341, the protein was designed to form a monomer. So, residues at the dimeric interface (the N- and C-termini and the hairpin turn) all gained new contacts, changing the environments of these residues (Fig. S6). However, as shown by the overall RMSD, these changes did not affect the overall structure of each monomeric subunit.

We also examined the local stability of designs HEEH_TK_rd5_0958 and HEEH_TK_rd5_0341 by hydrogen deuterium exchange (HDX) NMR. The HDX opening free energies revealed differences in local stability in different regions of the topology. The most stable secondary structure was Helix 2 for both miniproteins, with opening energies around 4 kcal/mol at 15 °C (compared to ~2-3 kcal/mol in Helix 1). The central β-hairpin was the least stable structure in HEEH_TK_rd5_0341 (Fig. 3J). Four residues in this hairpin (I21, G23, I24, and V26) form intramolecular hydrogen bonds that should protect those amides from exchange (Fig. S7A) but three of these residues exchanged too quickly in HEEH_TK_rd5_0341 to be measured by NMR. In contrast, three of the four hairpin residues that form intramolecular hydrogen bonds in HEEH_TK_rd5_0958 had measurable protection from exchange (Fig. 3G, Fig. S7B) and were similarly stable to Helix 1. Overall, the hierarchy of stabilities between Helix 2, Helix 1, and the central β-hairpin suggests the folding energy landscape is not fully cooperative.

The highest opening energy in the monomeric HEEH_TK_rd5_0958 was 4.5 kcal/mol, observed at I35 (Figs. 3J, G). This highest opening energy typically indicates the global stability of the protein (26), making HEEH_TK_rd5_0958 almost 2 kcal/mol more stable than the previous highest stability observed for a designed αββα structure (1). However, this higher stability was observed at a lower temperature (15 °C instead of 25 °C in ref. (1)) and in the presence of D_2_O, which typically stabilizes proteins.

### Stability determinants of αββα designs from a restricted design strategy

We next investigated which design features correlated with folding stability. To this end, we computed over a thousand structural and sequence-based metrics for each design and analyzed whether particular metrics correlated with stability. Several of the strongest individual correlations are shown in Fig. 2. Designs were generally more stable if their Rosetta energy scores were lower (Fig. 2B) and had more hydrophobic residues and hydrophobic sidechain contacts (Fig. 2C-D). Hydrophobic residue count correlated more strongly with stability than Rosetta energy. Stability also increased if a design’s sequence was highly compatible with its local backbone structure (see Methods and Fig. 2E). Finally, increased net charge destabilized our designs, although the optimal net charge was slightly negative (Fig. 2F-G). This stability change was approximately linear with the square of the net charge, as expected (27).

We also explored whether specific residues could individually have large influences on the stabilities of the designs. Because all designs are based on an identical architecture, each position in the sequence shares an identical structural role in all designs. Using the binomial test, we identified positions where specific amino acid identities had large and significant changes on the success rates of the designs (Fig. S8). Two positions near the N- and C-termini stood out as particularly important. Positions 2 and 39 are near the tips of each helix and contact each other in space (Fig. S8D). Across the design set, leucine residues at these positions increased the success rate of the designs by 25-39%, whereas other residues such as glutamate and tryptophan decreased the success rate by similar amounts. These differences in success rates were highly significant (adjusted p-value < 10^-18^) (Fig. S8C). The importance of these residues suggests that termini of the helices play an especially important role in the overall stability of designed αββα miniproteins.

To further examine individual residue contributions to stability, we performed deep mutational scanning analyses (Fig. S9) on the six αββα designs whose structures we verified by CD (Fig. 3B). Using our protease sensitivity assay (Fig. S5C-D), we measured the folding stability changes for all single mutants of each design (Fig. S9). Four of the six mutational scans showed many destabilizing mutations from replacing nonpolar residues in both the helices and the strands. A fifth design (HEEH_TK_rd5_0420) showed a similar pattern, but the helical residues seemed less sensitive to mutations than the strands. The high stability score of HEEH_TK_rd5_0420 (at the peak of our assay’s dynamic range) may have limited us from resolving the stability effects of other mutations in the helices (Fig. S9C). The sixth design (HEEH_TK_rd5_0018) showed many destabilizing substitutions at nonpolar sites in the helices, but only a small number could be observed in the β-hairpin, suggesting the hairpin may be less structured in this design (Fig. S9A). Overall, the positions that were most sensitive to mutations (change in stability < 1) were found in the buried hydrophobic core (Figs. S10-11), in particular large hydrophobic residues (Fig. S11). In contrast, hydrophobic residues, as well as polar and charged residues, that were more solvent exposed in the design models were less sensitive to mutation (Fig. S11). The specific sequence-stability relationships shown in the mutational scanning data suggest that the designs fold into specific structures. Furthermore, the consistency between a nonpolar residue’s burial in the designed models and its sensitivity to mutation (Fig. S11) provides support that the stable designs fold into their designed structures.

Charged and polar residues also contributed to folding stability, although they were less important than buried hydrophobic residues. The top three polar positions that were most sensitive to mutation (average change in stability < −0.5) were positions 15 (end of first helix), 28 (helix-capping position) and 31 (start of second helix that forms hydrogen bonding with the backbone) (Fig. 3E, Fig. S12). These positions indicate the importance of polar interactions toward stabilizing our designs and also support that the designs fold into their specific designed structures.

However, our mutational data also revealed some unexpectedly stable mutants (Fig. S9). For example, we expected that mutants to G23 would be highly destabilizing because G23 should be critical for forming the central β-hairpin. However, in four of the six designs, mutants to G23 could actually increase folding stability (Fig. S9B-D, F). To investigate this, we predicted the structures of all mutant sequences using AlphaFold 2 (21). Although most mutants were predicted to have similar structures to the original design, some predictions (including mutants of G23) suggest the possibility of alternative, compact structures (Fig. S13).

### Modeling relative contributions of biophysical determinants on folding stability

Our previous analysis identified individual determinants of stability without considering how various features relate to each other. Hence, we next analyzed which protein features were the most important contributors to stability and how they compared to each other. Instead of prioritizing predictive accuracy, we used linear regression to build a parsimonious, interpretable, low-resolution model. Our moderately accurate model (r = 0.64, r^2^ = 0.41; Fig. 2H, Table S3) included ten features chosen for either their large individual contributions to stability or their biophysical interest. Adding all 25 additional Rosetta energy terms provides only a minimal improvement to this low-resolution model (Table S4).

To analyze the strengths of the different features, we compared the different coefficients both in terms of their importance within our dataset (e.g. the impact of a one standard deviation change in each term, Fig. 2I left) and in terms of their biophysical strength (e.g. the impact of one additional residue, contact, charge, etc., Fig. 2I right). By representing the features in these two ways, we were able to observe how each feature contributes to a design’s stability while holding all other features constant. Relative to the variance in the features, the count of large nonpolar residues is the largest contributor to folding stability (Fig. 2I). Additional biophysical determinants known to stabilize globular proteins (2, 28–30), such as contacts between adjacent nonpolar residues and Ser/Thr helix capping, contribute to folding stability as well (Fig. 2I). However, our model also points to the stabilizing role of nonpolar residues at the design ends, which is a feature specific to the αββα topology (Fig. 2I). Whereas previous studies on the relative importance of stability determinants were based on assays that changed one feature on individual proteins (31, 32), our large-scale testing enabled us to analyze over a thousand protein features on several thousand proteins in parallel. This, in turn, allowed us to develop a model that offers criteria for designing even more stable αββα miniproteins.

### Designing αββα miniproteins using a diversity-oriented design strategy

Our restricted-design strategy (Round 5) focused on improving the success rate of designing stable αββα miniproteins but at the cost of reducing their structural diversity. Because we were now able to successfully generate stable αββα designs, we next investigated whether we could loosen the design restrictions that we had imposed, increase the diversity of our αββα miniproteins, and identify additional determinants of stability. Hence, we designed a new round of “diversity-oriented” (Round 6) αββα miniproteins based on fourteen different protein architectures instead of one. This allowed designs to have a greater variety of helix, β-strand, and loop lengths, while keeping the overall size of the protein to 43 residues (Fig. 1B). In addition, we did not impose residue restrictions on β-strands or in the middle loop and permitted a greater number of hydrophobic residues.

Importantly, we used our Round 5 stability data to directly re-weight the Rosetta energy function. Using ridge regression, we adjusted the weights on the Rosetta energy terms to create the best correlation with our measured Round 5 αββα stabilities, while regularizing the regression to penalize large deviations from the original weights. With this approach, we created three new energy functions labeled “Minor,” “Medium,” and “Heavy” based on how much the weights deviated from the original weights. We used these three energy functions (and the original weights) to design our Round 6 designs (Fig. S13).

We generated ~ 20,000 designs and chose our final set of over five thousand αββα designs for experimental testing by identifying designs that had the greatest structural diversity, varied sequence identity (no closer than 28/43 residues), and an αββα topology as determined by the computer program PSIPRED (33). Notably, we prioritized structural diversity (Fig. S2) in our final selection instead of prioritizing the expected success rate. The median sequence identity across all pairs of sequences was 28% (42% if only nonpolar residues are considered) (Fig. S3). However, the diversity in amino acid composition (overall and nonpolar only) is lower than several known protein domains of similar sizes (Fig. S4). Each design is named HEEH_KT_rd6_####, in which KT indicates the designer (author K.T.), rd6 indicates these designs constitute a new “Round 6” following the previous rounds of αββα design, and #### is a design number.

### Stability determinants of αββα designs based on diversity-oriented design strategy

We tested the stabilities of our “diversity-oriented” αββα miniproteins (and matching scrambled sequences) using the high-throughput protease sensitivity assay (1). Surprisingly, 12% of our scrambled sequences had stability scores above 1, compared to 2% or fewer in previous rounds (Fig. 4A). We further found that scrambled sequences were most likely to be stable when they were very hydrophobic and when their sequences had high helical propensity as determined by DSSP (34, 35) (Fig. S14). This suggested that designed sequences might also be stabilized by these properties alone, even if they did not fold into their designed structures. To remove these potential “false positive” designs from our analysis, we restricted our analysis to designs with a lower nonpolar residue count and lower helical propensity (Fig. S14). Restricting our analysis in this way removed 25% of our total designs, while lowering the fraction of stable scrambles from 12% to 6% (Fig. 4B). The overall fraction of stable designs was 26% - still substantially above the “success rate” of the scrambled sequences (Fig. 4B).

**Fig. 4.**
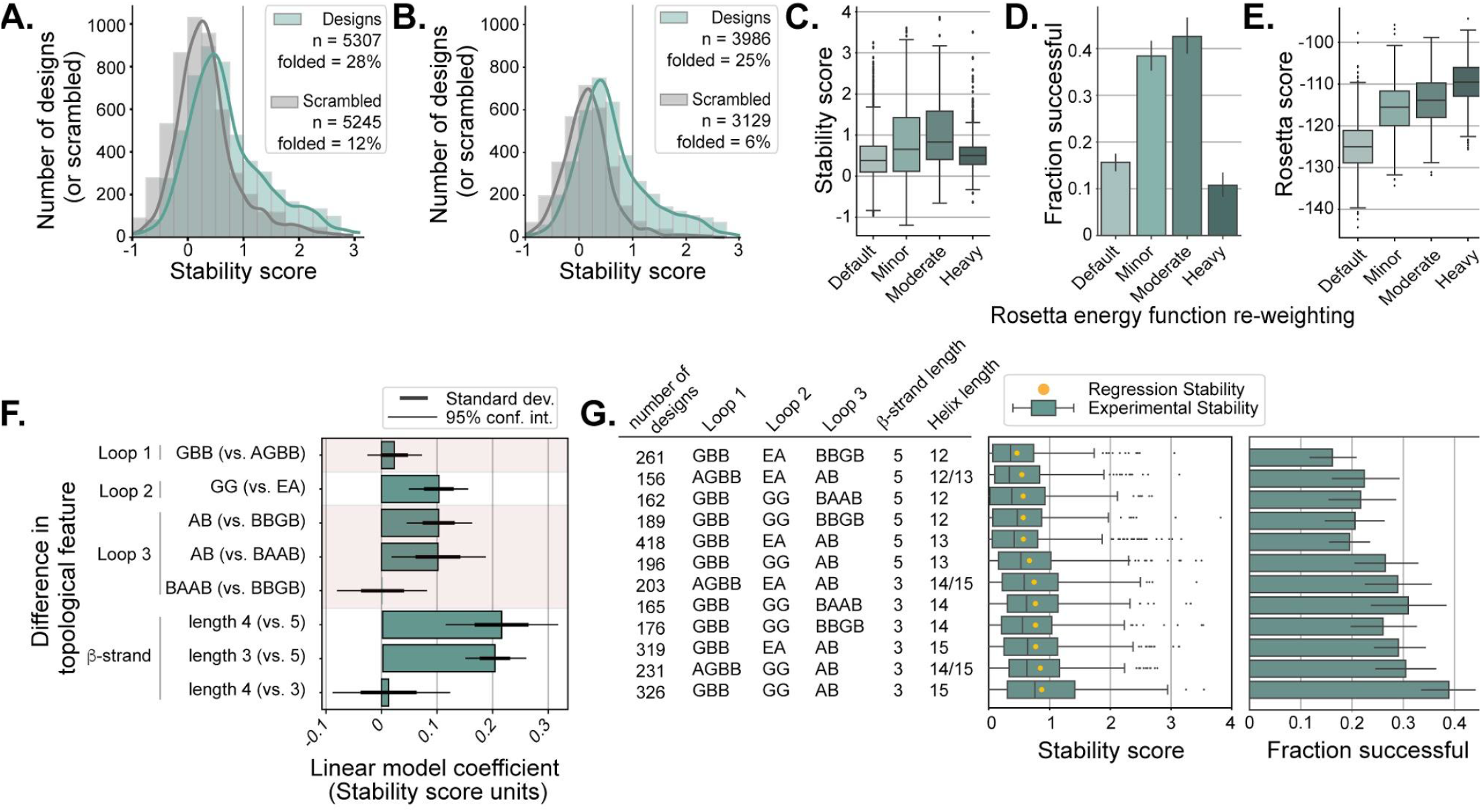
Experimental testing and analysis of αββα stability determinants from a diversity-oriented design strategy. (A) Stability score distribution of αββα miniproteins (green) and scrambled sequences (gray). (B) As in A, filtered to eliminate designed and scrambled sequences that may fold into nondesigned structures; see text and Fig. S7. The vertical line at stability score = 1 denotes the threshold above which we consider a design to be stable. (C-D) Stability scores and success frequencies of designs made with differently-weighted Rosetta energy functions; “Heavy” indicates the largest amount of reweighting. (E) Rosetta scores (using the unmodified score function) of designs made using different weighting; the more positive scores of the designs from the re-weighted energy functions indicate these designs are less favorable according to the default energy function. (F) Stability contribution of the most common loop patterns (using ABEGO notation) and β-strand lengths based on a linear regression model. (G) The most common unique structure combinations (loop pattern, β-strand and helix lengths) are listed (left) followed by the distribution of observed stability scores (middle, with the expected stability from the linear regression model as a yellow dot). At right, the fraction of stable designs for each unique structure. All error bars indicate 95% confidence intervals from bootstrapping.

We then analyzed the impact of differently weighted Rosetta energy functions on folding stability. On average, designs made using the reweighted energy functions had higher stability than designs made with the default energy function (Fig. 4C-D). However, some regularization (restraining the weights near their original values) was critical to successful re-weighting: the “Heavy” energy function, where the changes to the weights were the largest, performed much more poorly than the energy functions with “Minor” and “Moderate” changes to the weights (Fig. 4C-D). The success of the re-weighted energy functions suggests that empirical re-weighting could be an efficient practical tool for protein design in situations where large-scale data is available for a specific task. The designs created by the re-weighted energy functions would not have been favored under our previous design procedure, with larger changes to the weights leading to designs that appear less and less favorable according to the default energy function (Fig. 4E). These re-weighted “Minor” and “Moderate” energy functions also showed better correlation with previously published stabilities for other miniprotein topologies compared to the default score function (Table S5).

Next, we investigated how topological features (loop, β-strand, helix) of the designs affect folding stability. We selected the seven most common loop structures found in our designs (represented using ABEGO notation) (36) and the three most common β-strand lengths as inputs to another linear regression model (Fig. S15, Fig. 4F). The explanatory strength of this model is weak (95% conf. int. from bootstrapping, mean r = 0.167, mean R^2^ = 0.028). This is due to the simplicity of the model and because the topology-only model excludes critical stability determinants such as hydrophobic residue count. Despite these shortcomings, this model still enables us to examine the relative importance of different topological components. The largest structural contributors to stability are the lengths of β-strands and helices, with shorter β-strands (and corresponding longer helices) as the most favorable topological parameter (β-strand and helix lengths are inversely related because all designs have a fixed length of 43 residues) (Fig. 4F). Secondarily, particular structures in loops 2 and 3 influenced folding stability as well. A loop structure of GBB in the first loop, GG in the second loop, and AB in the third loop increases the stability of a design more than other loop structures (Fig. 4F).

Based on this topology-focused model, we would expect αββα miniproteins with a GBB-GG-AB loop patterning, β-strands that are 4 residues long, and helices that are 14-residues long to be more stable on average than αββα miniproteins with any other loop, strand, and helix combination (Fig. 4F). Although designs with a β-strand length of 4 residues were not common in our dataset, a very similar design structure (GBB-GG-AB with a β-strand length of 3 residues) had the highest average stability score and the highest success rate in our dataset (Fig. 4G), which is in agreement with a previous study on loop patterning and stability (37). In fact, this design pattern is the protein architecture that we used to generate all the Round 5 αββα miniproteins (Fig. S1A). However, the high success of this architecture in Round 6 may be due to using re-weighted energy functions that were optimized based on Round 5 designs with this specific architecture. Nonetheless, when we subset our Round 6 designs to identify αββα miniproteins with a GBB-GG-AB loop pattern and features that we previously determined to promote stability, these designs are diverse in their sequence identity and highly stable (81% successful) (Fig. 5). This provides a “recipe” for designing new stable αββα miniproteins in the future.

**Fig 5.**
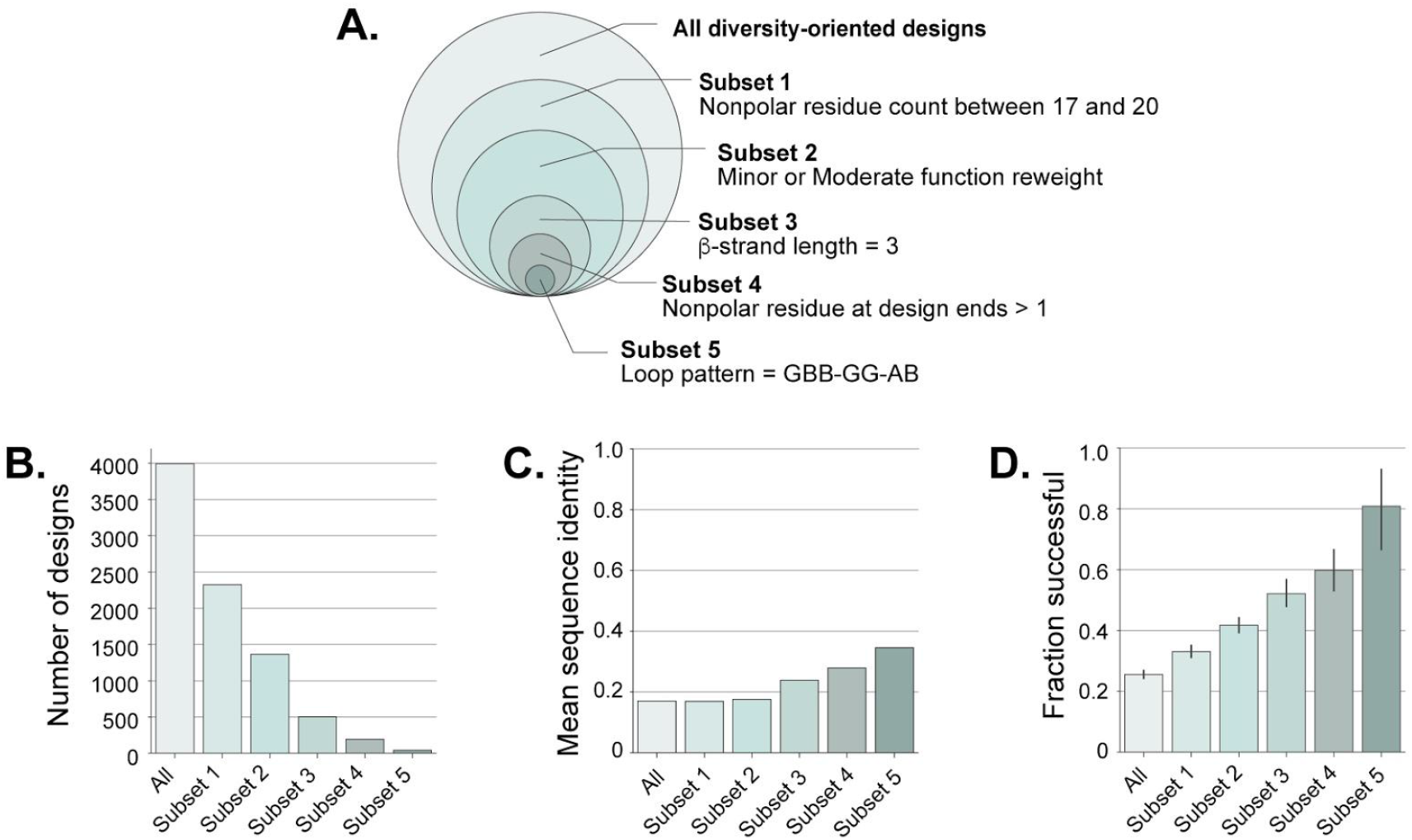
A recipe for building diverse high-stability αββα designs. (A) Designs made from a diversity-oriented strategy are grouped into subsets based on five features that we identified to be important for stability (Fig. 2I, Fig. 4F). (B) The number of designs that comprise each subset; (C) the mean sequence identity between any two designs in each subset; (D) the fraction of successful designs in each subset, with error bars indicating 95% confidence intervals from bootstrapping. Ideal designs (those with the parameters of Subset 5) are 80% successful with under 40% sequence identity between pairs of designs.

### Predicting stable de novo αββα miniproteins by AlphaFold 2

When we designed and tested αββα miniproteins for their folding stability, AlphaFold 2 was not yet available. With its recent release (21), we wondered whether AlphaFold 2 could discriminate between stable and unstable miniproteins. We explored this possibility even though AlphaFold 2 is intended for structure prediction and not stability prediction. Out of the ~5,600 and ~4,000 restricted and diversity-oriented designs, respectively, we found that 78% of the former and 20% of the latter had at least one predicted structure within 2□ RMSD to the designed model. These predictions were equally in agreement with design models regardless of whether a design was experimentally unstable, moderately stable, or stable, indicating that AlphaFold 2 did not discriminate stable from unstable designs (Fig. S16). We also examined whether the Rosetta energy scores of the AlphaFold 2-predicted models were better correlated with experimental stability scores than the scores of the original design models. The AlphaFold 2-predicted models did not improve the correlation with experiment for the Round 5 design set, but provided a small improvement for Round 6 (Fig. S17A-D). Neither RMSD nor AlphaFold 2’s average confidence measure (pLDDT) showed much ability to enrich for stable designs (Fig. S17E), indicating that AlphaFold 2 is currently unable to determine the folding stability of these designed miniproteins.

## Discussion

Understanding the biophysical determinants that enable proteins to fold and remain stable is important in protein design, drug development, and other areas. Here, we examined the stability determinants of the αββα miniprotein fold, which was previously identified as unusually challenging to design (1). We took advantage of an improved Rosetta design protocol (23, 24) to design over ten thousand αββα miniproteins using both restrictive and diversity-oriented design strategies. Our two design strategies led to over three thousand new stable designs (~2,100 restricted and ~1,000 diversity-oriented designs) and a much higher success rate (38%, Fig. 2A) than the 2% success previously reported (1). Our designed proteins also had a much higher success rate than control sequences with identical residue composition and polar-nonpolar patterning. This suggests that their stability was conferred by their designed three-dimensional structures. Supporting this, NMR structures of two designs closely matched the designed models (below 2 Å backbone RMSD, Fig. 3), circular dichroism spectra of six designs were consistent with the designed structures (Fig. 3C), and deep mutational scanning analysis of 5/6 designs showed specific sequence-stability relationships that were consistent with the designed structures. However, the lower resolution of circular dichroism and mutational scanning cannot directly demonstrate the atomic accuracy of the designs.

Our large dataset of stable designs enabled us to quantify determinants of stability for the previously-challenging αββα fold (Figs. 2I, 4F-G). Most of the stability determinants were common across globular proteins (2, 28, 29, 37, 32, 38–40), and similar to those previously observed in large-scale *de novo* design experiments (1). We also identified that designing hydrophobic residues near the termini was especially important for the αββα miniprotein fold (Fig. 2I). Our design success rate improved substantially when we used our large dataset to re-weight the Rosetta energy function specifically for αββα design (Fig. 4C-E). These observations largely explain the low success of previously designed αββα proteins: previous designs frequently employed non-optimal loop patterns, helix capping residues, and residues near the design termini, and typically had 13-16 nonpolar residues rather than the 17-20 used here (Fig. S1). Notably, the total number of nonpolar residues in each design is influenced by the design energy function and by parameters that restrict the amino acids that are sampled at each position according to the solvent accessibility of that position (41, 1). These restrictions are manually tuned to balance stability and solubility, as well as to reduce the search space of sequences. Designing proteins with too few nonpolar residues can thus be considered a failure of manual tuning as well as a failure of the design energy function.

Our study has several notable limitations. First, some fraction of “stable” designs are likely stable for non-designed reasons, such as folding into an alternative structure, forming a compact “molten” state, or aggregating on the surface of yeast. In our diversity-oriented set, 6% of our scrambled sequences met our stability threshold, compared with 25% of designs (Fig. 4B). Naively, this suggests that one in five stable designs could be stable for non-designed reasons. In addition, three of the six designs based on the restrictive protocol exhibited some oligomeric species when purified from *E. coli* (Fig. 3B), suggesting designs might be stabilized by intermolecular interactions. Because our regression analysis assumes that each design’s stability (or lack thereof) is due to its designed monomeric structure, our analysis will be unreliable if non-designed structures or interactions played an important role in our observed stabilities. Still, our regression analysis was able to identify specific three-dimensional features as stabilizing or destabilizing, such as buried unsatisfied polar atoms and attractive or repulsive ion pairs (Fig. 2I).

Secondly, our findings regarding the determinants of stability are limited to the specific context we examined: αββα miniproteins designed by a particular computational procedure. The samples of designs that we tested were not random: they were designed to be high stability and showed variation across some dimensions but not others. If a biophysical property (such as backbone torsional strain or higher polarity) varied only minimally across our design set, we would not be able to identify the contribution of that feature to stability. An alternative design procedure might also generate structures in a different region of “property space,” permitting high stability designs that are different from the recipe described in Fig. 5. Constructing a fully general model of folding stability will ultimately require a broad sampling of sequences, structures, and biophysical properties. Our work here investigating a specific design space suggests that this should be possible.

Despite these limitations, our study demonstrates how large-scale experimental testing can be applied to solve a challenging design problem and to quantify the biophysical features that influence design stability. In contrast to other studies that use mutagenesis to study determinants of folding stability (19, 42–44), our method examines the strengths of different biophysical features across thousands of different protein contexts, although these contexts are all related by the αββα fold and design procedure. Simplified low-resolution models like our linear regression are valuable for building biophysical intuition about the strengths of different interactions (45, 46) as well as for guiding the construction of high-resolution models like the Rosetta energy function, which is also an additive model (47). Our stable αββα designs (and our recipe for generating more) may also be valuable scaffolds for engineering binding functionality for therapeutic, diagnostic, and synthetic biology applications (13, 48, 49).

## Materials and Methods

### Computational protein design

We designed αββα miniproteins using Rosetta based on our previous work (1). Briefly, we used fragment assembly to build backbones according to protein architectures specified in a blueprint file (22), as in (41, 1). For the restricted design strategy (Round 5), we chose the protein architecture that previously led to generating the greatest number of stable (defined only here as stability score ≥ 0.8) αββα miniproteins (Fig. S1A). This architecture restricted the αββα miniprotein structure to have two helices that are 14-residues long, two β-strands that are 3-residues long, and three loops with an ABEGO pattern of GBB, GG, and AB, respectively. We also applied several design constraints. We forced the first residue in the middle loop (position 22) to be nonpolar (AFILMVWY), the solvent-facing residues in the β-strands (positions 20 and 25) to be polar or charged (QNSTDEHKR), and any helical positions were prevented from being designed as Gly, Thr, or Val as they are known to destabilize helix formation (50–52). We also required all designs to possess at least 15 hydrophobic residues (AFILMVWY) and no more than 21 hydrophobic residues. Finally, we filtered out designs with low Rosetta total energy scores or low buried nonpolar surface area (Fig. S1B-D).

Sequence design was performed using the Rosetta protocol FastDesign (24), the beta_nov16_protease version of the full-atom energy function, and a recently-improved sampling method designed to prevent over-compaction (53). In order to select αββα miniproteins for experimental testing, we ranked each αββα design by their predicted stability scores, which was determined by a Lasso regression model that we built using previous αββα miniprotein structural metrics and experimental stability scores (1). Based on this ranking, we selected the top ~5,600 designs with a threshold of 67% sequence identity for experimental testing.

Round 6 designs were designed as above with several changes. First, we utilized 15 different protein architectures. Moreover, we removed the hydrophobic restriction in the middle loop, were more permissive on non-helical residues (GDNST) inside the helices, and allowed hydrophobic residues to appear on the protein surface. We further specified a penalty for a protein’s net charge outside the range of −5 or 3. Upon generating ~20,000 αββα designs, we took several steps to select over 7,000 designs for experimental testing. We first built Lasso and XGBoost regression models (54) using experimental data from Round 5 to identify ~3,000 designs with significantly different predictions between the two models (predicted stability scores were at least 0.25 scores away from the best-fit line between the models). We next independently performed principal component analysis to identify ~9,000 designs that were most distant from each other. From the combined ~12,000 αββα designs, we selected ~7,400 designs for experimental testing whose sequence identity was no closer than 66% to any other design.

Although all ~7,400 designs were experimentally tested, we determined afterwards that many of these structures either diverged away from the αββα topology during design or were not predicted to fold into an αββα structure by Rosetta’s ab initio algorithm. We further found that scrambled sequences could form secondary structure according to psipred (33). To focus our analysis on designed αββα structures, we restricted our analysis to ~5,300 designs meeting these criteria: distance between the C-terminus to the middle loop < 22 ⍰, distance between the N- and C-termini < 20 ⍰, β-strand lengths according to DSSP ≤5 residues, loop lengths ≤ 5 residues, and unbroken αββα secondary structural elements according to DSSP (34, 35).

### Energy function re-weighting

The Rosetta energy function is a weighted sum of individual, independent score terms (47). To test whether our experimental data could directly optimize the energy function for αββα miniprotein design, we sought to re-weight these terms in Round 6 to produce the best correlation with our experimentally measured stability scores from Round 5. In re-weighting, we also sought to bias our new weights to be as close to the original weights as possible by using ridge regression (54). However, because the L2 regularization in ridge regression biases coefficients to be near zero, we used ridge regression to identify optimal *perturbations* to our original weights, rather than directly optimizing the weights themselves. To determine the appropriate perturbations, we first regressed our set of experimentally measured stability scores against the original Rosetta (computational) total scores of the designs. We then used the residuals from this regression (i.e. the error in the prediction of experimental stability score) as the target values for our ridge regression. We used scikit-learn’s implementation of Ridge regression (54) to determine new weights on the 25 unweighted Rosetta score terms that best fit the residuals of the first regression (Fig. S7). The coefficients in this second regression are effectively perturbations to the original Rosetta weights that minimize the error in predicting experimental stability scores (subject to the regularization constraint). After performing ridge regression, the new score function weights were determined based on the formula:

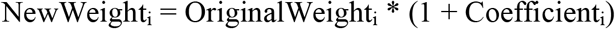

where NewWeight_i_ is the new weight on score term *i*, OriginalWeight_i_ is the original weight for term *i* in the beta_nov16_protease energy function, and Coefficient_i_ is the coefficient on score term *i* in the ridge regression.

We tested three new weight sets (“Minor,” “Medium”, and “Heavy”) in addition to the default weights. These new weight sets were determined using three different regularization strengths in the ridge regression and are named based on the magnitude of the change. The “Minor” set used regularization strength alpha=200,000; “Moderate” used alpha=20,000, and “Heavy” used alpha=0.1. “Heavy” corresponds to the value of alpha from a cross-validated ridge regression using scikit-learn’s RidgeCV method (54). In all weight sets, the score term fa_intra_rep_xover4 was maintained at its default value to avoid favoring extended structures. These weight sets are all provided in the supporting information.

### Miniprotein library generation

We reverse translated the residue sequences and optimized the codons (based on *E. coli* codon frequencies) of all αββα miniproteins that we selected for experimental testing using DNAworks 2.0 (55). We also included scrambled sequences (while preserving locations of P and G residues as well as nonpolar/polar patterning) for each corresponding αββα sequence (following (56)). Both oligo libraries (Round 5, and Round 6 + mutational scanning) were purchased from Agilent.

### Yeast display and protease stability assay

DNA amplification, yeast display proteolysis, sorting, and next-generation sequencing were all performed by research contract to the University of Washington BioFab (57) according to the protocol of (1). Yeast display was performed using a display vector with improved protease resistance (24).

### Computing stability scores

We calculated a “stability score” for each design based on a probabilistic model described previously (1). The model determines the EC_50_ (the protease concentration at which half of the yeast cells pass selection during flow cytometry) for each design. The difference between the experimental EC_50_ in the folded state and predicted EC_50_ in the unfolded state (based on the identical model from (1)) is what we call a “stability score.” The overall stability score for each sequence is the minimum of the independent stability scores measured by trypsin and chymotrypsin. As previously (1), Data were filtered based on the confidence interval of the EC_50_ estimate: only sequences where the 95% confidence interval was smaller than 2.0 (meaning the equivalent of two selection rounds, or 9x protease concentration) were retained for analysis. However, the mutational scanning data were not filtered based on the EC_50_ confidence interval; however, only 6/4650 (0.1%) sequences had low confidence stability estimates.

### Computing metrics and Regression modeling

The Rosetta models used for computing structural features were the lowest energy structures from at least 1,000 ab initio trajectories and 200 relax trajectories starting from the design models. Rosetta design models were scored using the Rosetta score function, and we computed structural and biophysical features pertaining to secondary structure, dipeptides, hydrophobicity, hydrogen-bonding, and fragment quality using the score_monomeric_designs package (https://github.com/Haddox/score_monomeric_designs).

For regression modeling, we performed linear regression by bootstrapping (sampling with replacement 1000 times) using Python scikit learn (54) and selected the 95% confidence interval for each variable’s coefficient for analysis. For the restricted design strategy, we first used stepwise linear regression to identify eight features (large nonpolar count, nonpolar residue-residue contacts, local sequence-structure propensity, Ser/Thr at helix caps, Glu-Arg residue-residue contacts, nonpolar residue at design ends (which we define as positions 1, 2, 42, and 43 of the 43-residue-long protein structure), Glu-Glu residueresidue contacts, and increased net charge) that increased the correlation coefficient between predicted and experienced stability scores. We also selected two features (favorable net charge at helix ends and buried unsatisfied polar atoms) to determine their relative contributions to stability. For the diversity-oriented topology-focused linear regression model, we selected the seven most common loop patterns and the three most common β-strand lengths found in our dataset as inputs to a linear regression model.

### Calculation of local sequence-structure agreement

The compatibility of each protein sequence with its local backbone structure (Fig. 2E) was computed using the abego_res_profile method from (1).

### Protein expression and purification

We purchased six αββα designs whose nucleotide sequences were optimized for *E. coli* expression and encoded in the pET-28a(+) vector (that contains an N-terminal His-tag and thrombin cleavage) from Twist Bioscience. The plasmid vectors were transformed in BL21(DE3) competent cells (Invitrogen or Sigma Aldrich) and grown overnight in a starter culture of 50 mL LB media (Fisher Bioreagents) and 50 μg/mL kanamycin at 37°C while shaking at 225 rpm. 16-18 hrs later, we inoculated 500 mL of LB media and 50 μg/mL kanamycin with 10 mL of the starter culture and allowed the competent cells to grow until OD_600_ ~0.6-0.8.

In preparation for NMR analysis, we transformed one αββα design encoded in pET-28a(+) into BL21(DE3) competent cells (Sigma Aldrich) and grown in an LB media starter culture (as stated above). After 16-18 hrs, we pelleted the cells by centrifugation, replaced the LB media with M9 media (40 mM Na_2_HPO_4_, 8.5 mM NaCl, 20 mM KH_2_PO_4_, 60 mM d-Biotin, 55 mM Thiamine, 0.1 mM CaCl_2_, 0.01 mM ZnSO_4_, 2 mM MgSO_4_, 50 ug/mL kanamycin) that included 15 mM ^15^NH_4_Cl and 10 mM ^13^C glucose (Cambridge Isotopes) and resuspended the pellet. We then inoculated 500 mL of LB media with M9 media (including 15 mM ^15^NH_4_Cl, 10 mM ^13^C glucose and 50 μg/mL kanamycin) with 10 mL of the resuspended starter culture and allowed the competent cells to grow until OD_600_ ~0.6-0.8.

Afterwards, for both labeled labeled and unlabeled competent cells, we induced protein expression by adding a final concentration of 500 mM Isopropyl β-D-1-thiogalactopyranoside (IPTG) (Fisher Bioreagents) to the LB media and allowing the cells to grow overnight at 15°C while shaking at 225 rpm. We then harvested the cells by centrifugation at 4°C and lysed the cells in 30 mL of lysis buffer (20 mM Tris, 500 mM NaCl, 30mM imidazole, 0.25% CHAPS, 1mM PMSF, pH 8.0), which included 60 mg of chicken lysozyme (Sigma), 1.5 μL of benzonase nuclease (Sigma Millipore), and 1 tablet of Pierce protease Inhibitor EDTA-free (ThermoFisher) followed by sonication (QSonica SL-18).

Next, we separated insoluble bacterial material by centrifugation (10,000 x g for 30 min) and purified the αββα miniproteins by immobilized metal-affinity chromatography (IMAC), which involved transferring the supernatant onto Econo-pac columns (Bio-Rad) that were previously prepared with Ni-NTA (Qiagen), washing the column with 15 mL Wash Buffer (20 mM Tris, 500 mM NaCl, 30 mM imidazole, 0.25% CHAPS, 5% glycerol, pH 8.0), and eluting the samples in 10 mL of Elution Buffer (20 mM Tris, 300 mM NaCl, 500 mM imidazole, 5% glycerol, pH 8.0). We initially verified the size and purification of the miniproteins by SDS-PAGE electrophoresis and Coomassie stain gel analysis. Then, we concentrated them by a centrifugal filtration system (Amicon Ultracel-15 or Amicon Ultra-0.5).

We further purified both labeled and unlabeled miniproteins by size-exclusion chromatography (Bio-Rad NGC Chromatography System) using a Superdex 75 10/300 GL column (GE Healthcare) and eluted in PBS buffer. A mixture of proteins with known molecular weights (BSA, Ovalbumin, Ribonuclease A, Aprotinin, and Vitamin B12) were also separated by SEC to determine a calibration curve, which was used to designate expected miniprotein monomeric and dimeric fractions (see Fig. 3B). Miniprotein size and purification were verified first by Coomassie stain gel analysis and then by mass spectrometry (Synapt G2 Si, Waters).

### Circular dichroism

Far-ultraviolet circular dichroism measurements were performed on six αββα designs (HEEH_rd5_0018, HEEH_rd5_0341, HEEH_rd5_0420, HEEH_rd5_0614, HEEH_rd5_0958, and HEEH_rd5_3711) using a Jasco J-815 spectrophotometer. All analysis was performed on the unmodified expression constructs including a 21-residue N-terminal linker (MGSSHHHHHHSSGLVPRGSHM). We measured the concentration samples with a Qubit 4 Fluorometer (Invitrogen) and diluted them to a final concentration of ~0.1-0.4 mg/mL in PBS buffer. Wavelength scan measurements were made using a 1 mm path-length cuvette from 195 to 260 nm at 25°C and 95°C. We also measured temperature melts at 220 nm for every 1°C from 25°C to 95°C. For temperature melt analysis, we smoothed the data with a Savitsky-Golay filter of polyorder = 3.

### Nuclear Magnetic Resonance

NMR spectra for HEEH_TK_rd5_0341 and HEEH_TK_rd5_0958 structure calculations were acquired at 288 K, on Bruker spectrometers operating at 600 or 800 MHz, equipped with TCI cryoprobes with the protein buffered in 20 mM sodium phosphate (pH 7.5, 150 mM NaCl) at concentrations of ~ 0.5 to 1.0 mM. Resonance assignments were determined for ^15^N/^13^C-labeled protein using FMCGUI (58) based on a standard suite of 3D triple and double-resonance NMR experiments collected as described previously (59). All 3D spectra were acquired with non-uniform sampling in the indirect dimensions and were reconstructed by the multi-dimensional decomposition software qMDD (60), interfaced with NMRPipe (61). Peak picking was performed manually using NMRFAM-Sparky (62). Torsion angle restraints were derived from TALOS+ (63). Automated NOE assignments and structure calculations were performed using CYANA 2.1 (64). However, for HEEH_TK_rd5_0341 NOEs identified manually with high confidence, that were not consistent with a monomeric structure, were added as initial restraints for dimeric structure calculation. The best 20 of 100 CYANA-calculated structures were refined with CNSSOLVE (65) by performing a short restrained molecular dynamics simulation in explicit solvent (66). The final 20 refined structures comprise the NMR ensemble. Structure quality scores were performed using Procheck analysis (67) and the PSVS server (68). Longitudinal (T1) and transverse (T2) 15N relaxation rates were determined for the constructs at 288 K (at 600 MHz) (69) using the integrated signal from the structured amide regions (>8.5 ppm) of 1D 15N-edited spectra, fitted to an exponential decay as a function of delay times. Rotational correlation times (τc) were estimated based on T1-T2 ratios (70) and hydrodynamic radii (rH) calculated using the Stokes-Einstein equation.

### Hydrogen-deuterium exchange and analysis

NH/D amide exchange rates were determined for HEEH_TK_rd5_0341 and HEEH_TK_rd5_0958 (including the 21aa N-terminal linker MGSSHHHHHHSSGLVPRGSHM) by performing an exchange series at 288 K (at 600 MHz), by monitoring the decay rate of amide peak intensities in ^1^H-^15^N HSQC spectra collected over the course of 24 hrs. HEEH_TK_rd5_0958 dissolved in MES buffer at ~ 500 μM (pH 5.8, 150 mM NaCl) was lyophilized, and exchange was initiated by solubilizing in an equal volume of D_2_O; each HSQC time point was acquired in ~ 5 minutes. For HEEH_TK_rd5_0341, exchange was initiated by mixing phosphate buffered protein (at ~ 1.0 mM, pH 7.5) with D_2_O at a ratio of 1:19, and each HSQC time point was acquired in ~ 16 minutes. The first time points for the series were started ~ 5 minutes following the initiation of exchange. Peak intensities were fitted to a single exponential decay (with an offset due to the presence of residual 5% H_2_O for HEEH_TK_rd5_0341). Opening free energies were calculated from these rates as previously described (71–73). For residues where exchange was too fast to quantify, we calculated an “upper limit” ΔG_open_ based on an exchange rate of 0.1 min^-1^ (the fastest quantifiable rate was 0.066 min^-1^).

### AlphaFold analysis

All αββα miniprotein structures were predicted from their primary sequences using AlphaFold 2 (21) without using multiple sequence alignment (MSA) information because αββα miniproteins have low similarity to natural proteins. Five models were generated for each sequence and the lowest RMSD model to the designed structure was used for the analysis in Figs. S13, S17, and S18.

### Analysis of residue composition difference

The residue composition of all Round 5 and 6 were compared by taking all possible combinations of protein pairs, and first identifying the total number of unique residues for each protein. Then, we took the absolute difference for each unique residue, taking the sum of all absolute differences, and then dividing the sum by the total number of residues being compared. Other protein domains were compared, LysM (https://pfam.xfam.org/family/LysM), PASTA (https://pfam.xfam.org/family/PF03793), and Cold Shock (https://pfam.xfam.org/family/CSD).

### Analysis of structural agreement among Rosetta design, AlphaFold, and NMR models

Different models were aligned to each other using PyMOL. To analyze the RMSD between the Rosetta design model and AlphaFold model, both structures were aligned (super structure1, structure & name n+c+ca+o). To analyze the RMSD between the Rosetta design model (or AlphaFold model) with the NMR ensemble, all twenty states of the NMR ensemble were aligned to each other, and the design model (or AlphaFold model) was added to the NMR pdb file as the 21st state. Then, RMSD values for all NMR states to the design model (or AlphaFold model) were determined (intra_fit structure////n+c+ca+o, 21), and the average RMSD determined (Fig 3F, H).

## Supporting Information

All blueprint files, Rosetta input commands, design files, and datasets are provided in https://github.com/tekim1/abba_protein_stability_manuscript.

## Acknowledgements

This work was supported, in part, by the National Institute of General Medical Sciences through award number 1DP2GM140927 and award number 5T32GM105538. K.T. was supported by JSPS KAKENHI grant number (19J30003) and is currently supported by a Human Frontier Science Program Long-Term Fellowship. Computational work was supported in part through the resources and staff contributions provided for the Quest high performance computing facility at Northwestern University (which is jointly supported by the Office of the Provost, the Office for Research, and Northwestern University Information Technology). NMR work was performed by the Structural Genomics Consortium, a registered charity (no: 1097737) that receives funds from Bayer AG, Boehringer Ingelheim, Bristol Myers Squibb, Genentech, Genome Canada through Ontario Genomics Institute [OGI-196], EU/EFPIA/OICR/McGill/KTH/Diamond Innovative Medicines Initiative 2 Joint Undertaking [EUbOPEN grant 875510], Janssen, Merck KGaA (aka EMD in Canada and US), Pfizer and Takeda. Yeast display selections and next generation sequencing were performed by the University of Washington BioFab. CD spectroscopy was performed using Northwestern University’s Keck Biophysics Facility. We further thank the members of the Rocklin lab for discussions and comments on this manuscript.

