## supplemental figures and tables for "Dissecting the stability determinants of a challenging de novo protein fold using massively parallel design and experimentation"

[1] Driskill Graduate Program in Life Sciences, Northwestern University Feinberg School of Medicine, Chicago, IL 60611
[2] Department of Pharmacology, Northwestern University Feinberg School of Medicine, Chicago, IL 60611
[3] Center for Synthetic Biology, Northwestern University, Evanston, IL 60208
[4] Structural Genomics Consortium, University of Toronto, Toronto, ON, M5G 1L7, Canada.
[5] Princess Margaret Cancer Centre and Department of Medical Biophysics, University of Toronto, Toronto, ON, M5G 2M9, Canada.
[6] Chemistry for Life Processes Institute, Northwestern University, Evanston, IL 60208
[7] Department of Biochemistry & Institute for Protein Design, University of Washington, Seattle, WA 98195
[8] Robert H. Lurie Comprehensive Cancer Center, Northwestern University, Chicago, IL 60611
[9] These authors contributed equally to this work

**Corresponding author:**
Gabriel J. Rocklin.


**Contents**:

Figures S1 to S
Tables S1 to S5
SI References

### Supplementary Figures

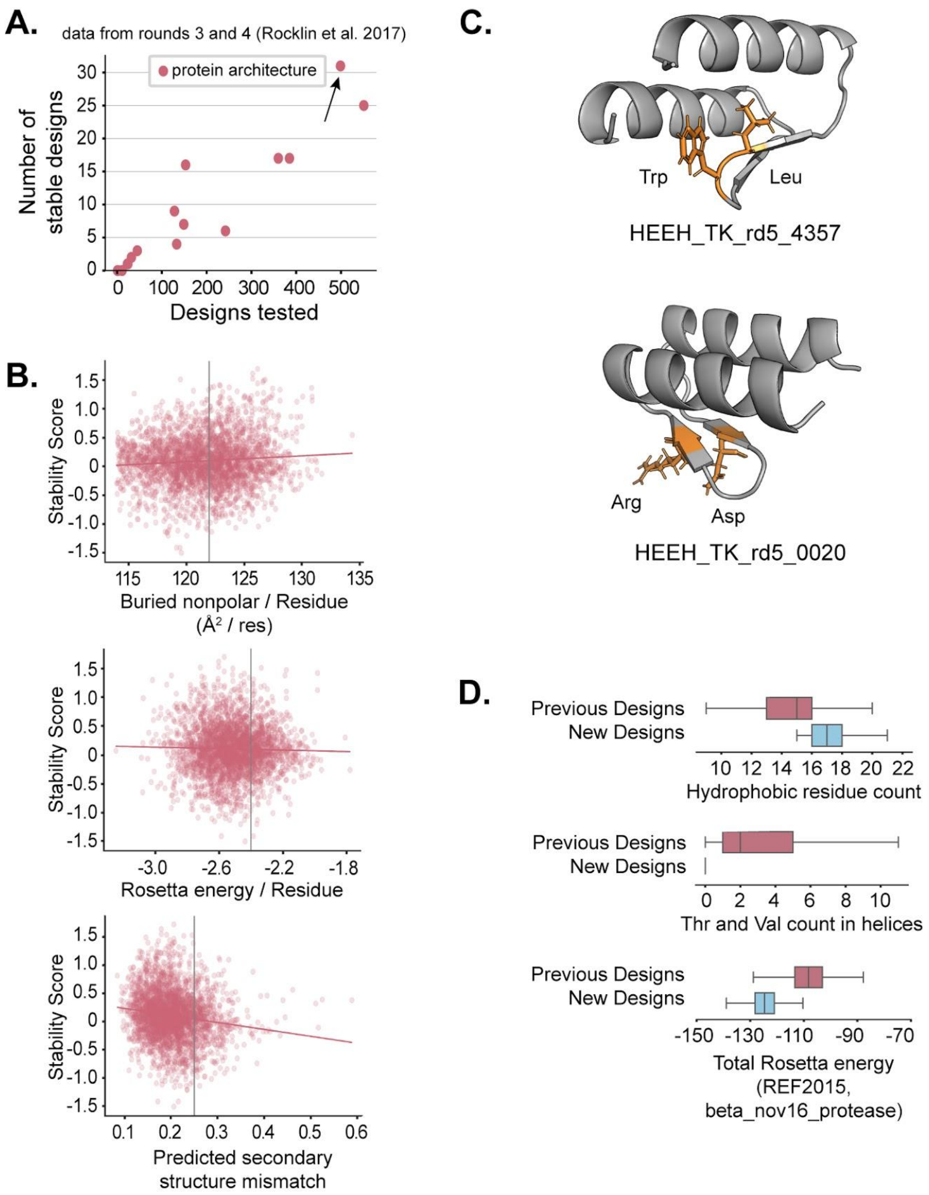


**Fig. S1. Restricted αββα miniprotein design strategy**. (A) We selected the protein architecture that led to generating the greatest number of stable designs (black arrow) from our previous study (1). Here, we set the threshold of stability as a design having a stability score ≥ 0.8 (B) Based on the Round 3 and 4 αββα design and stability data from our previous study (1), we set thresholds for specific features that a design can have: buried nonpolar surface area / residue > 122 (top), Rosetta energy / residue < -2.4 (middle), and predicted secondary structure mismatch < 0.25 (bottom); vertical gray line denotes the minimum or maximum threshold, and red line denotes best fit line. (C) We also forced all designs to have one hydrophobic residue in the middle loop (top) and polar/charged residues at solvent-facing β-strand positions (bottom); example residues are highlighted (orange) on a cartoon model. (D) We also set additional constraints to the αββα designs (blue) in comparison to our previous study (1): 15-21 nonpolar residues allowed (top) and no Thr/Val in helices (middle); using these restrictions, we were able to generate αββα designs whose Rosetta energies were lower than previously designed (bottom).

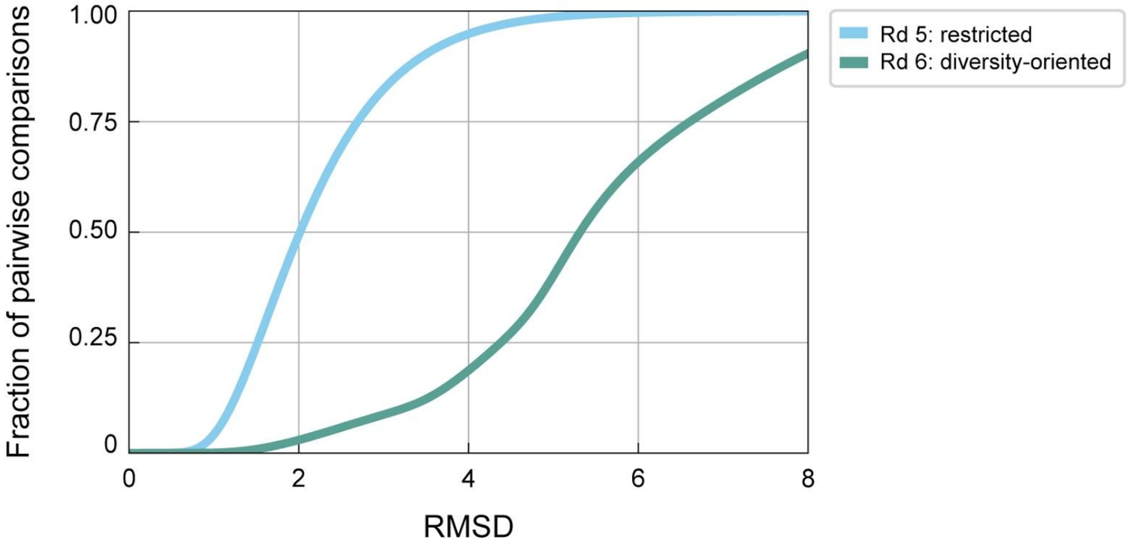


**Fig. S2. Structural diversity of restricted and diversity-oriented αββα miniprotein designs.** Each miniprotein design was compared to every design within the same library (Round 5 or Round 6) by the distance between alpha-carbons.

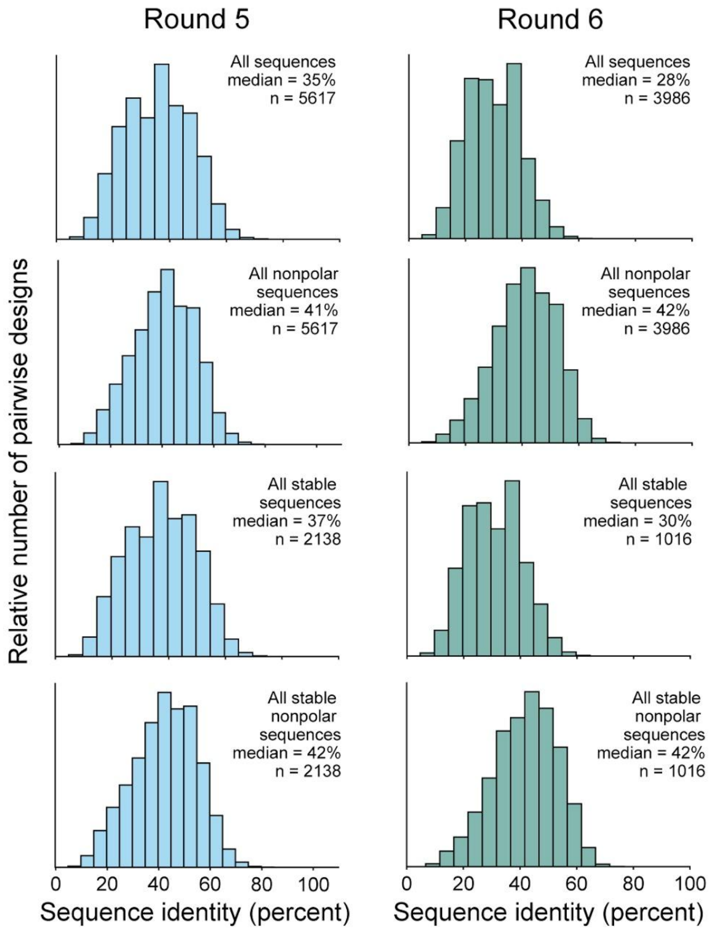


**Fig S3. Sequence identity of designs.** To evaluate the sequence diversity among Round 5 (restricted) and Round 6 (diversity-oriented) designs, we calculated the sequence identity (all sequences or all nonpolar sequences only) between all possible pairs of designs (all or stable). The distributions of the sequence identities are shown as histograms, with bins being 5 percentage points wide.

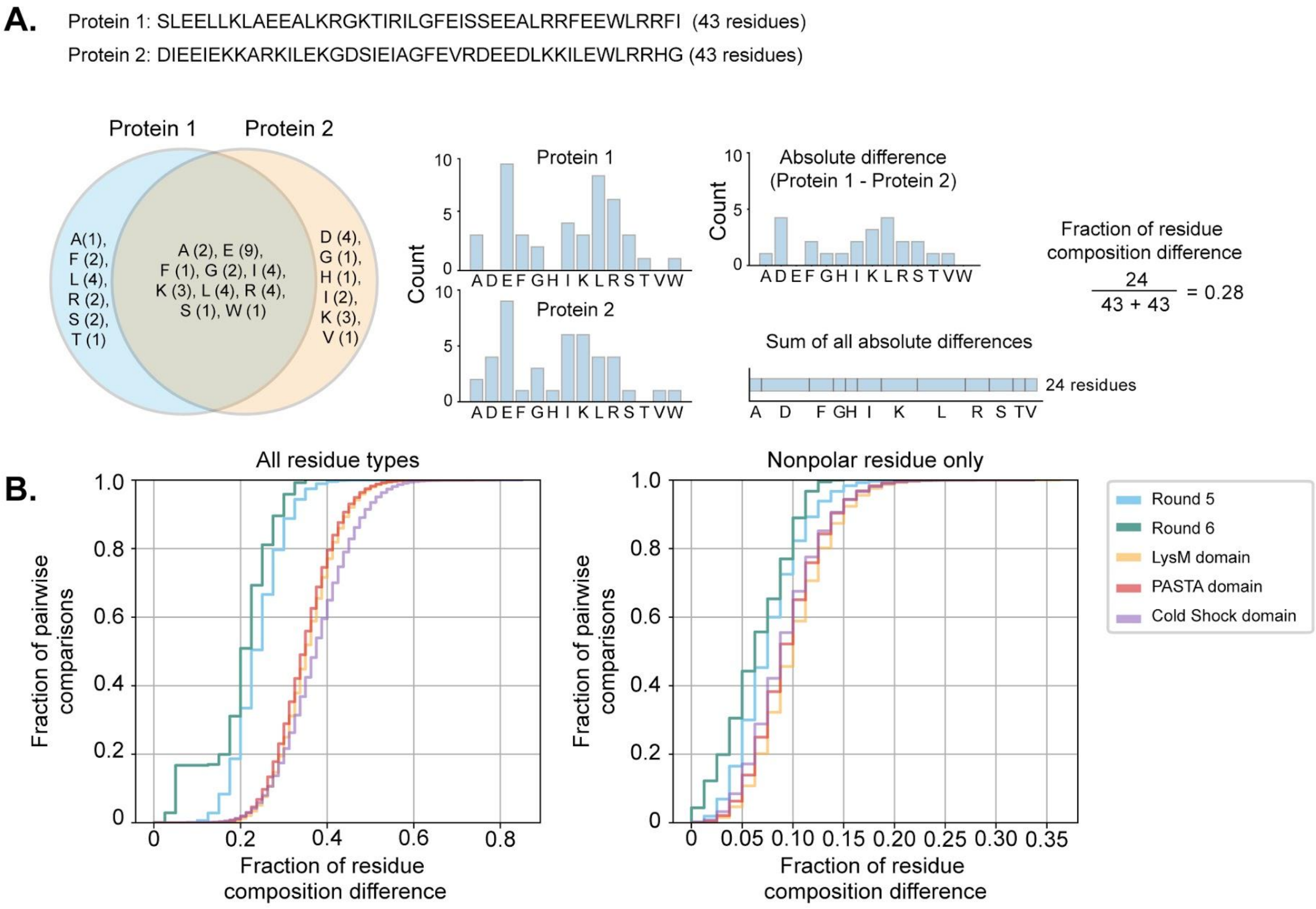


**Fig S4. Residue composition of αββα miniprotein designs in comparison to similarly-sized protein domains.** To compare the compositional diversity of αββα miniproteins with known protein domains of similar sizes (LysM: 44-65 residues (2), PASTA: ~70 residues (3), and Cold Shock: 65-70 residues (4)), we calculated (A) how different a pair of protein sequences are to each other by identifying the total number of unique residues for each protein sequence, taking the absolute difference for each number of unique residue, summing the absolute difference, and calculating the fraction of the sum over the total number of possible residues. (B) The fraction of pairwise comparisons for all possible protein pairs in Round 5, Round 6, and three natural protein domains are shown as a function of the fraction of residue difference (all residue types, left; nonpolar residues only, right).

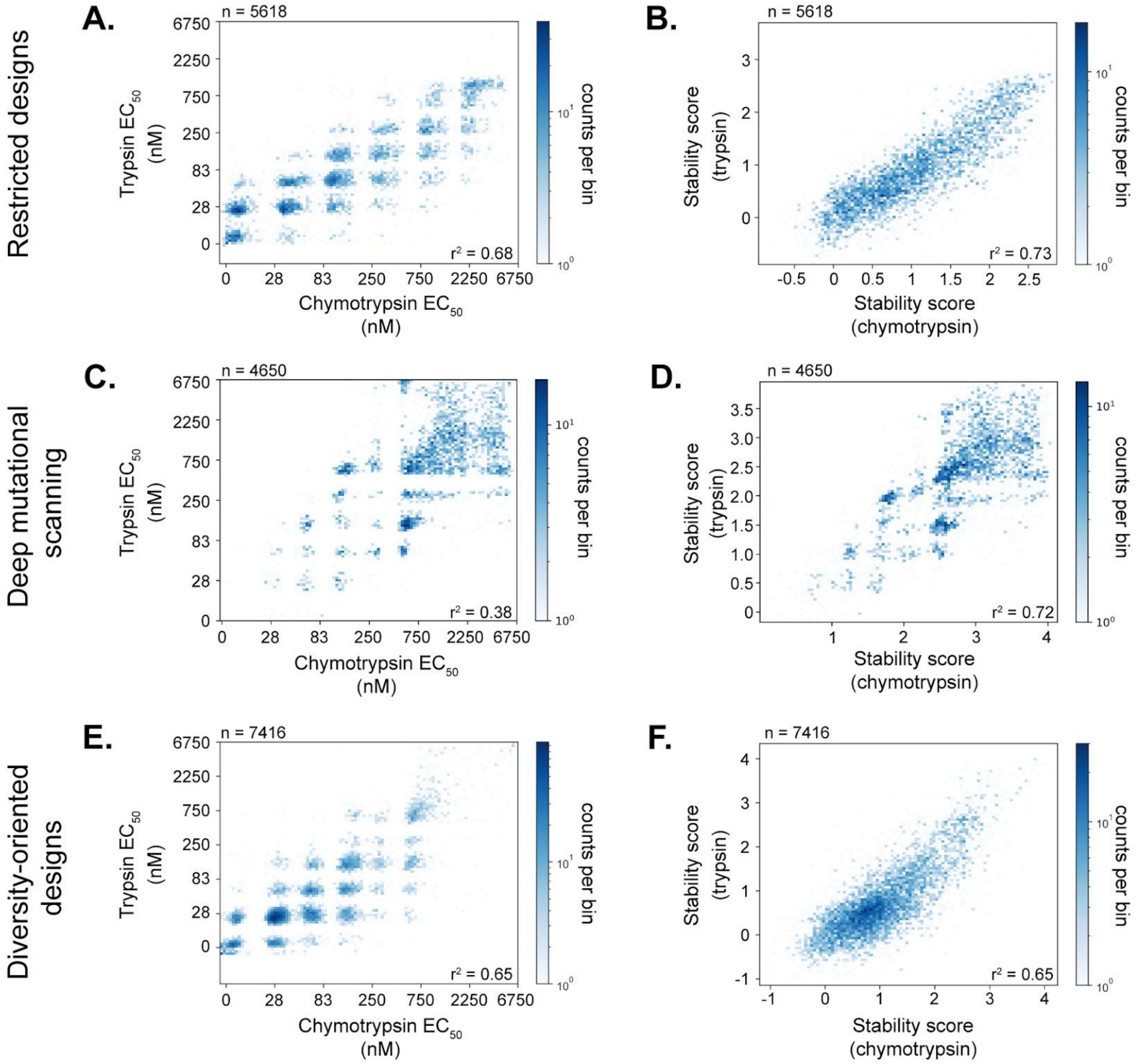


**Fig. S5. Protease stability assay using trypsin and chymotrypsin.** αββα miniproteins generated from (A-B) a restricted design strategy, (C-D) deep mutational scanning, and (E-F) a diversity-oriented strategy were displayed on the surface of yeast cells and subject to varying concentrations of either trypsin of chymotrypsin (from 0 to 6750 nM). $EC_{50}$ values (left) and calculated stability scores (right) at each protease concentration are depicted as 2D-histograms.

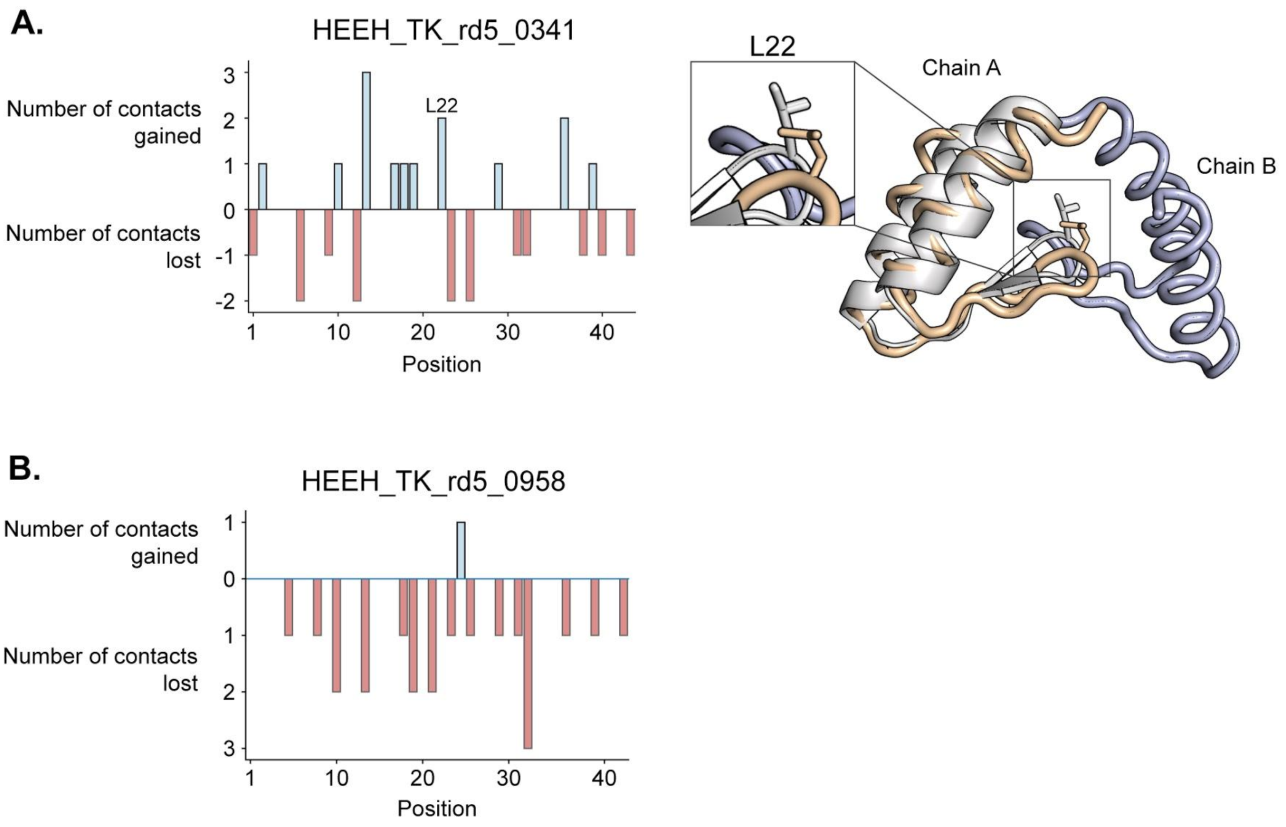


**Fig. S6. Structural agreement between Rosetta design model and NMR ensemble for HEEH_TK_rd5_0341 and HEEH_TK_rd5_0958.** (A-B) We determined the Euclidean distance between all possible residue-residue pairs (using the beta-carbon positions for each residue, excluding Gly) in the design model and an NMR model (whose beta-carbon positions after superposition were the average of an NMR ensemble consisting of twenty structures for each chain in HEEH_TK_rd5_0341 and twenty structures for HEEH_TK_rd5_0958). From the distance calculations, we quantified the number of contacts (< 8 Å) a residue in each position gained or lost from the design model to the NMR model. Cartoon of HEEH_TK_rd5_0341 design model (gray) is overlaid on Chain A (orange) of the NMR model.

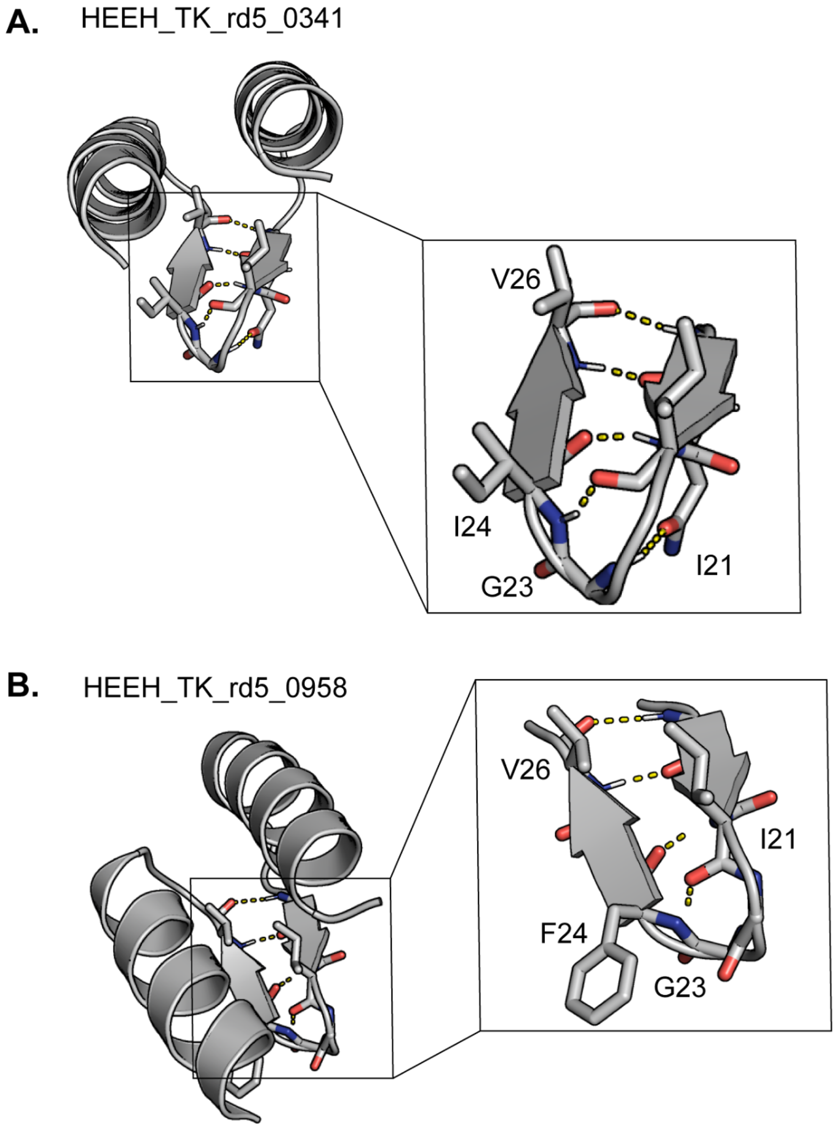


**Fig. S7. Intramolecular hydrogen bonds in the β-hairpin of HEEH_TK_rd5_0341 and HEEH_TK_rd5_0958.** Cartoons of (A) HEEH_TK_rd5_0341 and (B) HEEH_TK_rd5_0958 highlighting the hydrogen bonds (yellow) that are formed within the β-hairpin.

**A.**

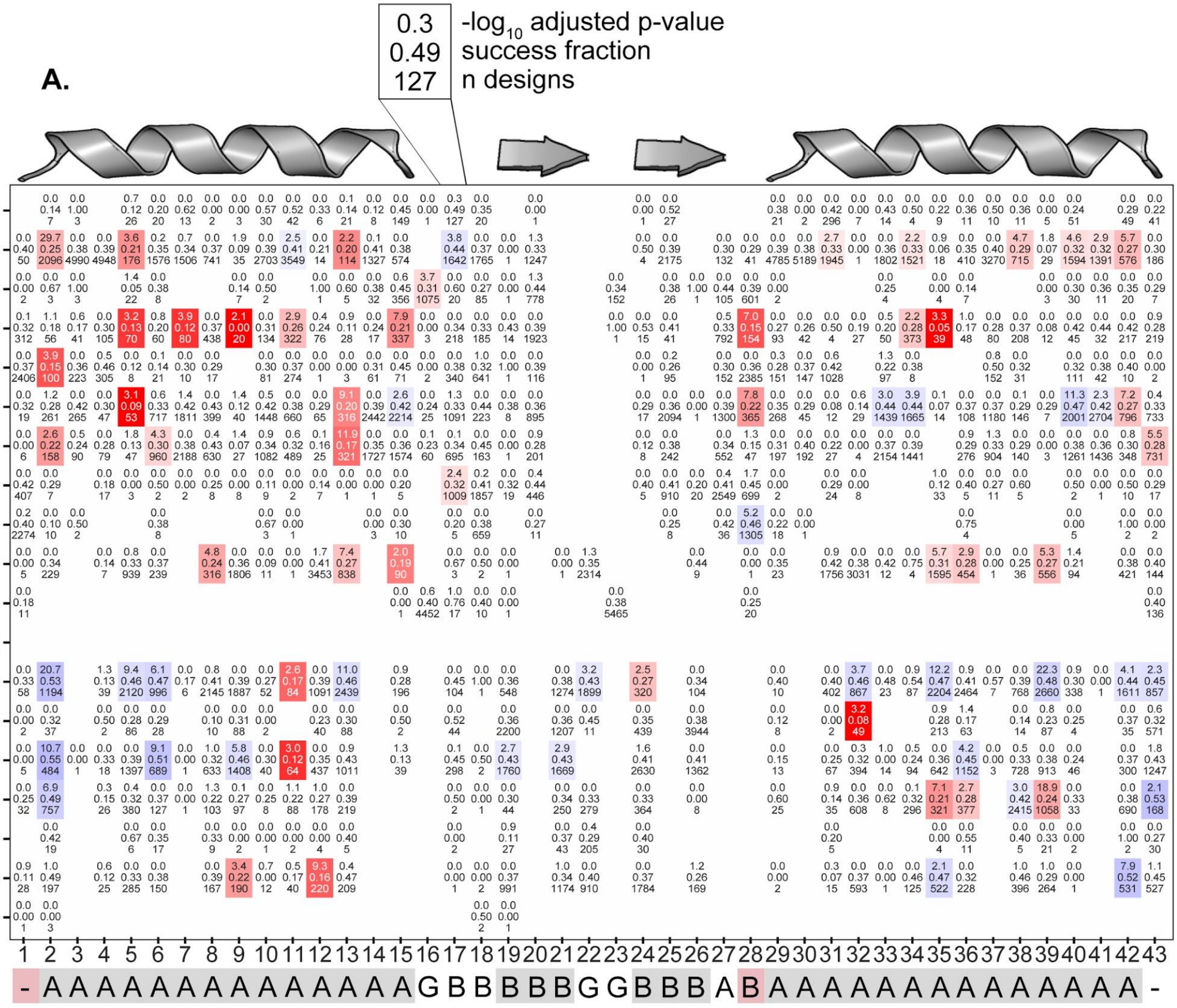

0.3
0.49
127
-log10 adjusted p-value
success fraction
n designs
Q
E
N
H
D
R
K
T
S
A
G
M
L
V
I
W
Y
F
P
1 2 3 4 5 6 7 8 9 10 11 12 13 14 15 16 17 18 19 20 21 22 23 24 25 26 27 28 29 30 31 32 33 34 35 36 37 38 39 40 41 42 43
- A A A A A A A A A A A A A A G B B B B B G G B B B A B A A A A A A A A A A A A A A -

**B.**

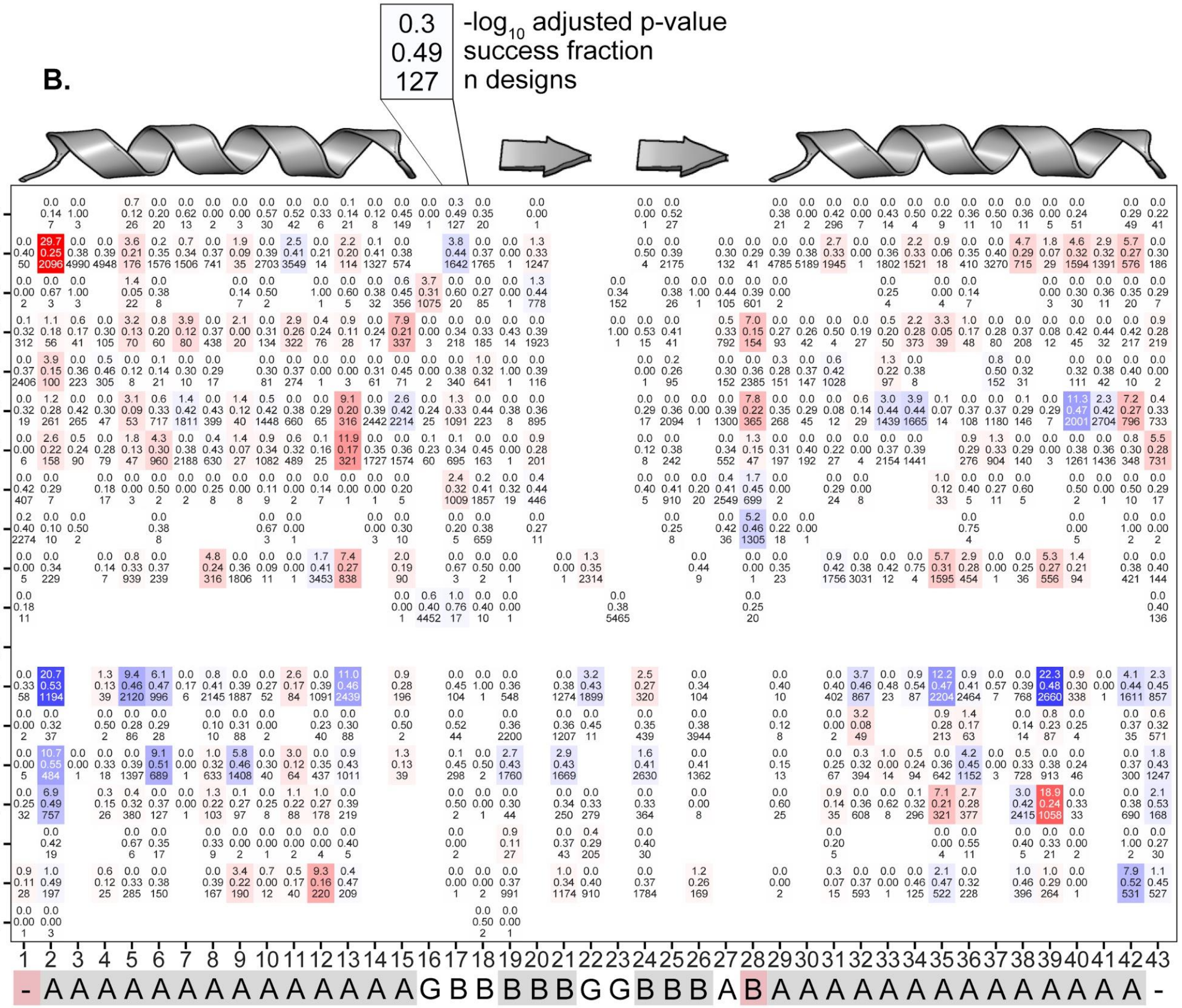

0.3
0.49
127
-log10 adjusted p-value
success fraction
n designs
Q E N H D R K T S A G M L V I W Y F P
1 2 3 4 5 6 7 8 9 10 11 12 13 14 15 16 17 18 19 20 21 22 23 24 25 26 27 28 29 30 31 32 33 34 35 36 37 38 39 40 41 42 43
- A A A A A A A A A A A A A A G B B B B B G G B B B A B A A A A A A A A A A A A A A -

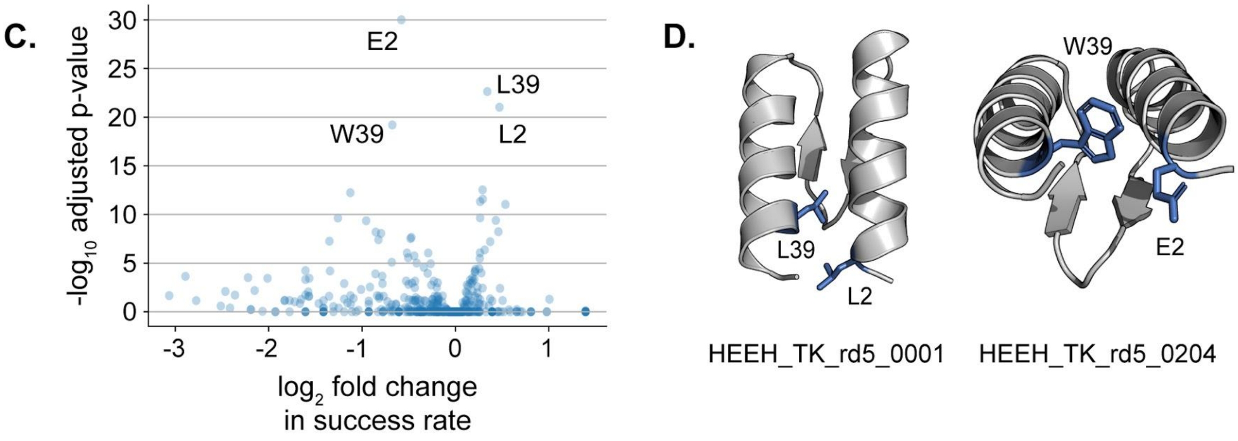


**Fig. S8. Contribution of specific residues on folding stability.** For αββα miniproteins made using a restricted design strategy (Round 5), we performed binomial tests for all possible residues in all 43 positions to examine whether the success rate of designs containing that residue differed from the overall success rate (0.38). All p-values were adjusted for multiple testing using the Benjamini–Yekutieli procedure as implemented in (5) (A-B) The p-value, success rate, and number of designs containing each amino acid at each position. In A, significant residues (adjusted p-value < 0.01) are colored according to the fold-change in success rate, with favorable amino acids in blue and unfavorable amino acids in red. In B, residues are colored by statistical significance. The ABEGO pattern of the Round 5 architecture is shown below; secondary structure is colored in gray and helix caps are shown in red. (C) Volcano plot indicating the p-value and change in success rate associated with each amino acid at each position. The most significant residue-positions are labeled on the plot (W39, E2, L39, L2). (D) Two representative αββα cartoons visualizing L2 and L39 (left) and E2 and W39 (right).

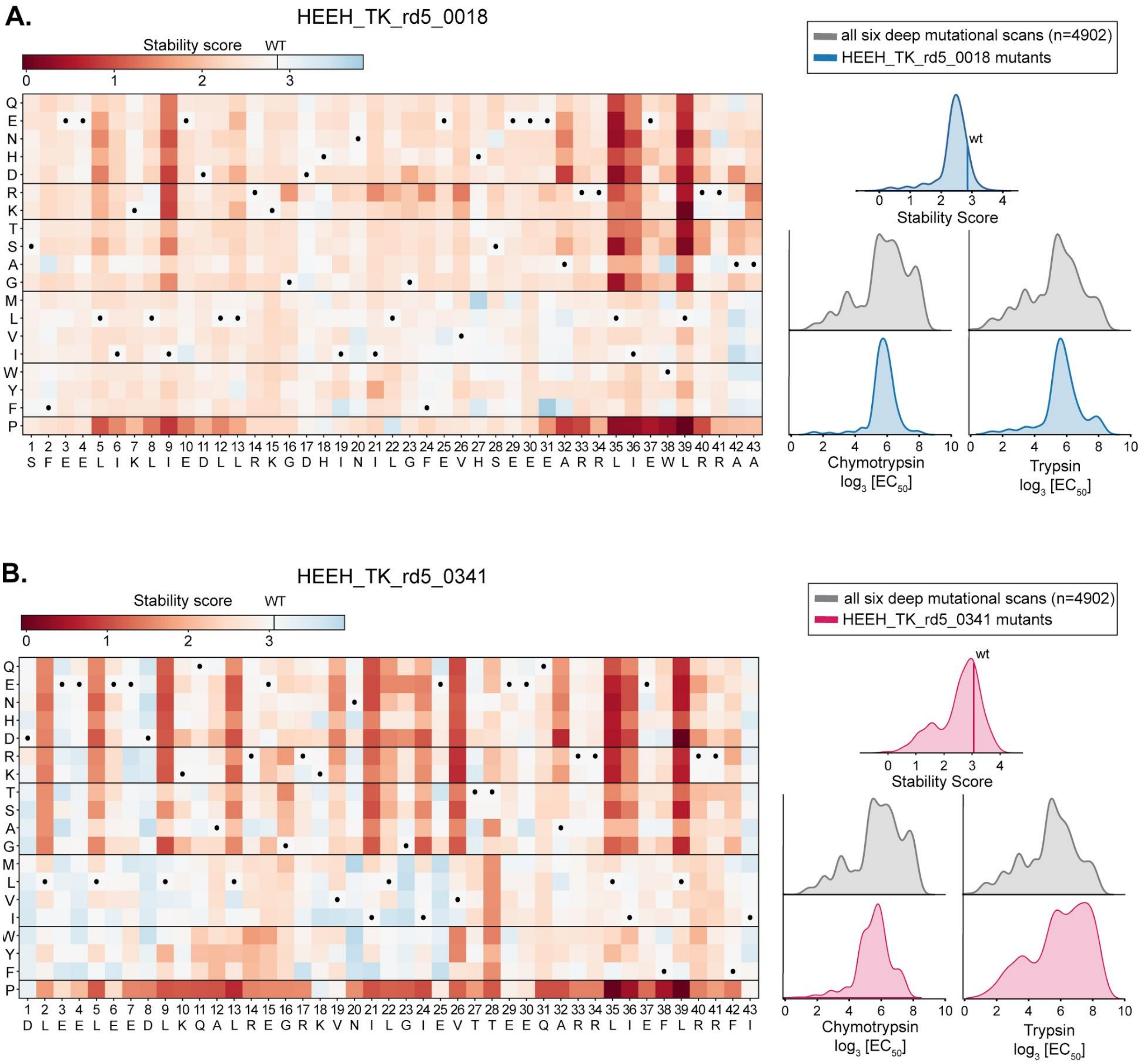

A.
HEEH_TK_rd5_0018
Stability score
WT
all six deep mutational scans (n=4902)
HEEH_TK_rd5_0018 mutants
wt
Stability Score
Chymotrypsin
log$_3$ [EC$_{50}$]
Trypsin
log$_3$ [EC$_{50}$]
S F E E L I K L I E D L L R K G D H I N I L G F E V H S E E E A R R L I E W L R R A A
B.
HEEH_TK_rd5_0341
Stability score
WT
all six deep mutational scans (n=4902)
HEEH_TK_rd5_0341 mutants
wt
Stability Score
Chymotrypsin
log$_3$ [EC$_{50}$]
Trypsin
log$_3$ [EC$_{50}$]
D L E E L E E D L K Q A L R E G R K V N I L G I E V T T E E Q A R R L I E F L R R F I

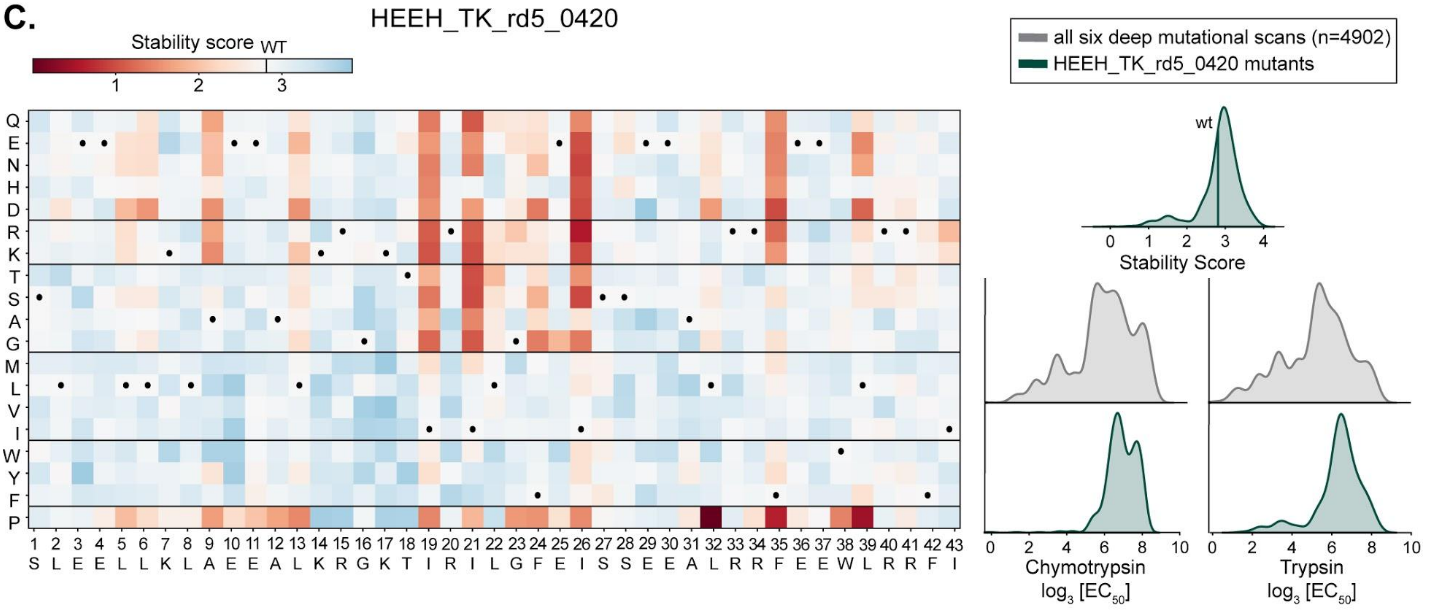
C.
HEEH_TK_rd5_0420
Stability score WT
1 2 3
Q E N H D R K T S A G M L V I W Y F P
1 2 3 4 5 6 7 8 9 10 11 12 13 14 15 16 17 18 19 20 21 22 23 24 25 26 27 28 29 30 31 32 33 34 35 36 37 38 39 40 41 42 43
S L E E L L K L A E E A L K R G K T I R I L G F E I S S E E A L R R F E E W L R R F I
all six deep mutational scans (n=4902)
HEEH_TK_rd5_0420 mutants
wt
0 1 2 3 4
Stability Score
0 2 4 6 8 10
Chymotrypsin
log3 [EC50]
0 2 4 6 8 10
Trypsin
log3 [EC50]

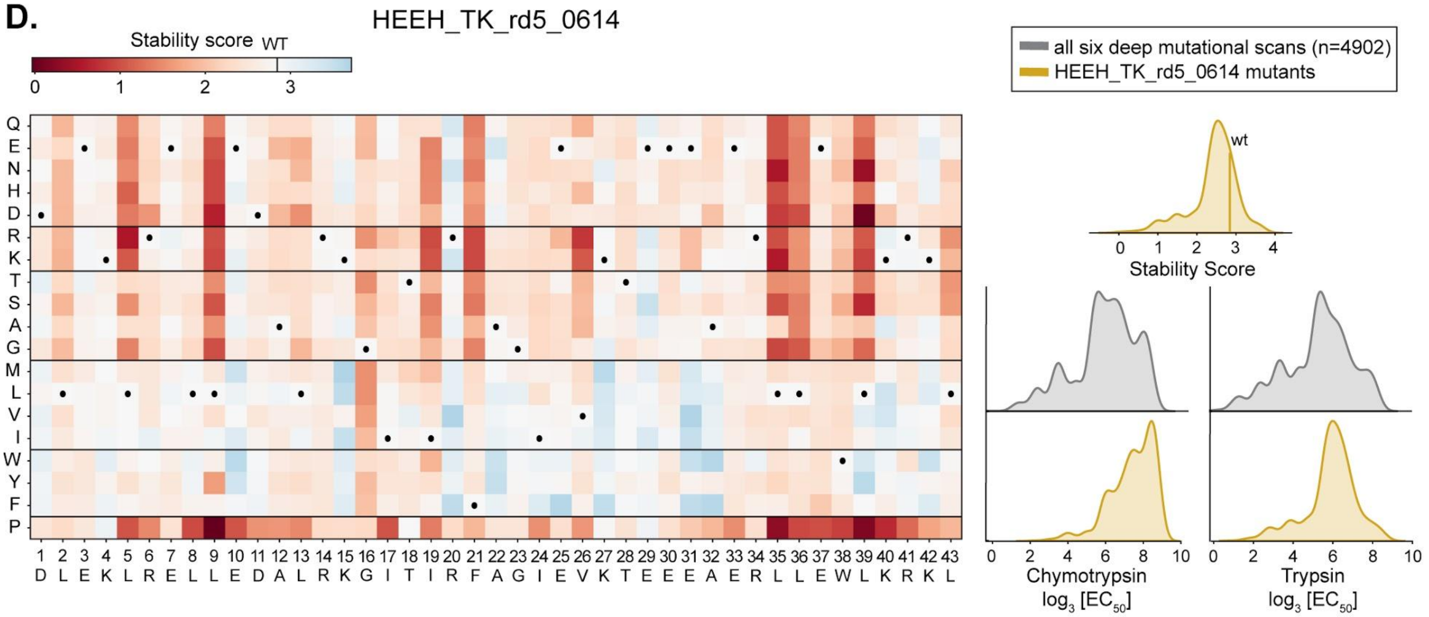
D.
HEEH_TK_rd5_0614
Stability score WT
0 1 2 3
Q E N H D R K T S A G M L V I W Y F P
1 2 3 4 5 6 7 8 9 10 11 12 13 14 15 16 17 18 19 20 21 22 23 24 25 26 27 28 29 30 31 32 33 34 35 36 37 38 39 40 41 42 43
D L E K L R E L L E D A L R K G I T I R F A G I E V K T E E E A E R L L E W L K R K L
all six deep mutational scans (n=4902)
HEEH_TK_rd5_0614 mutants
wt
0 1 2 3 4
Stability Score
0 2 4 6 8 10
Chymotrypsin
log3 [EC50]
0 2 4 6 8 10
Trypsin
log3 [EC50]

E.

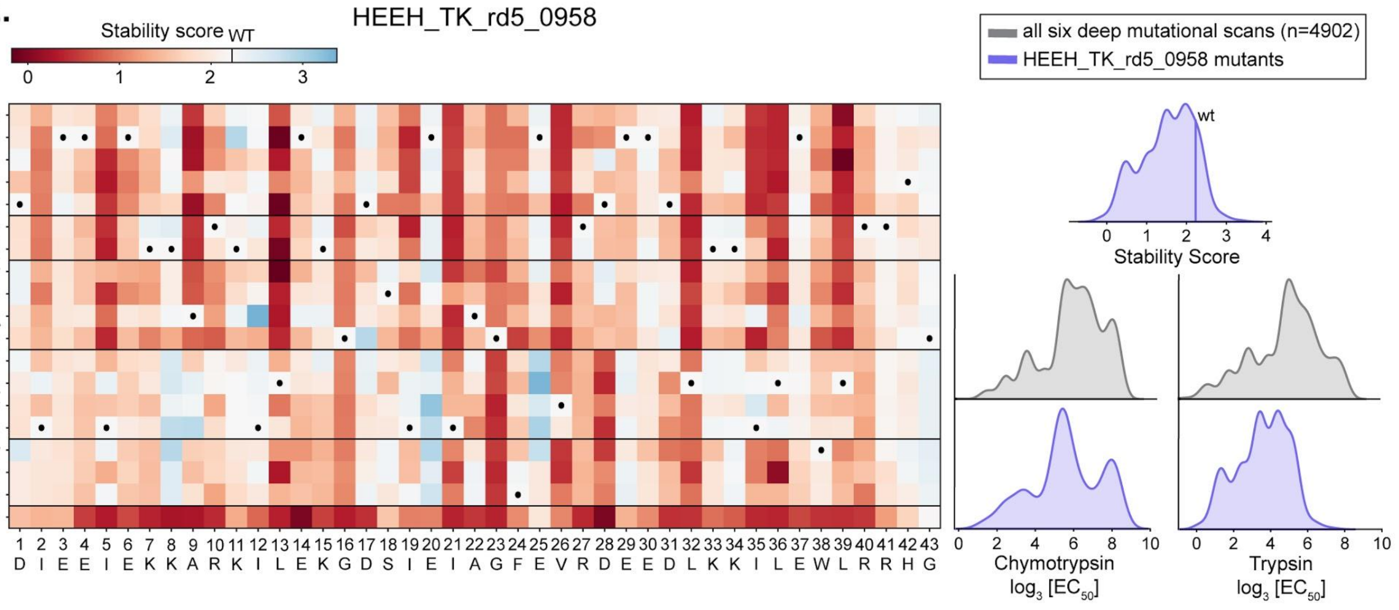


F.

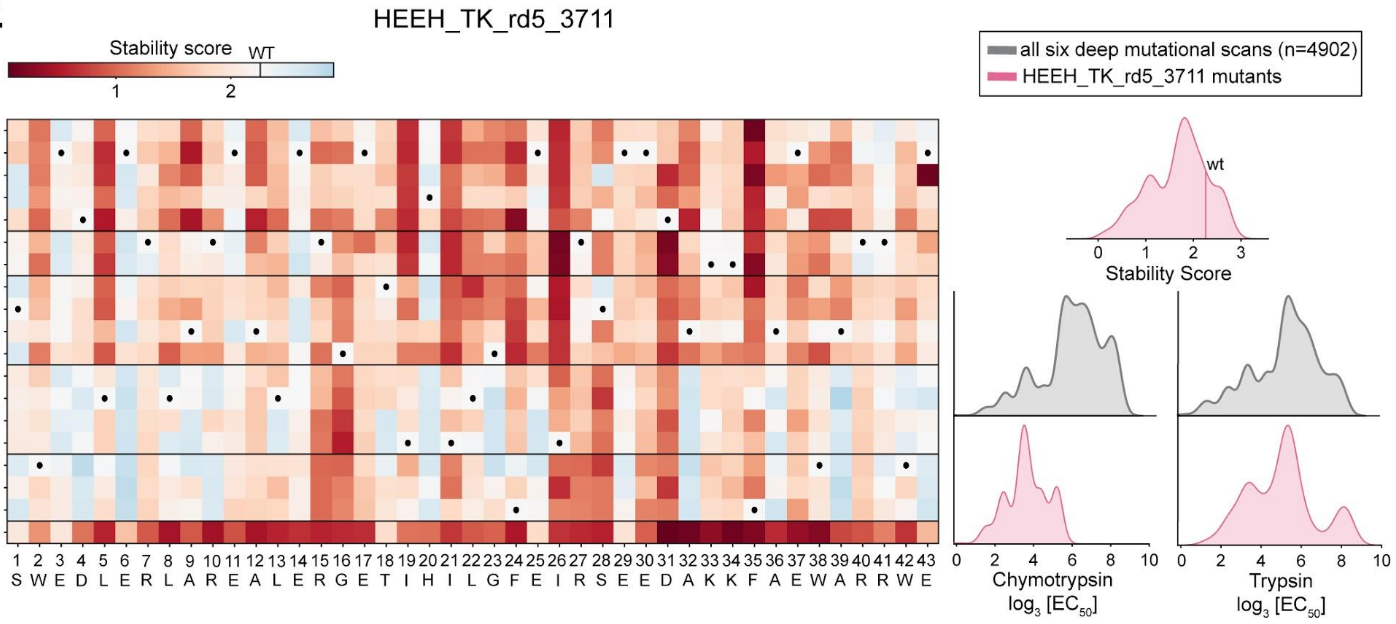


**Fig. S9. Stability scores of six αββα miniproteins by deep mutational scanning.** For six designs (Table 1) we created a library consisting of all possible single mutants and tested for their folding stability by a yeast display-based protease sensitivity assay (1). (Left) Results are shown as heatmaps. The wildtype residue is represented as a black dot, and the stability scores of each mutant relative to the wildtype is shown as a range from red (less stable than the wild type) to blue (more stable than wildtype). (Right, top) The distribution of stability scores for each design, (right, middle), $EC_{50}$ concentrations of protease for all six deep mutational scans, and (right, bottom) for each design are all shown as histograms. Mutants for five of the six designs were more likely to destabilize the protein when compared to their corresponding wild type (A-B, D-F). However, this is the opposite for HEEH_TK_rd5_0420 (C). Because the typical mutant in HEEH_TK_rd5_0420 does not seem to have particularly unique properties when compared to the other five designs, it may be that the high $EC_{50}$

values (e.g. for chymotrypsin) prevents us from discriminating the true differences between the wild type and mutant stabilities.

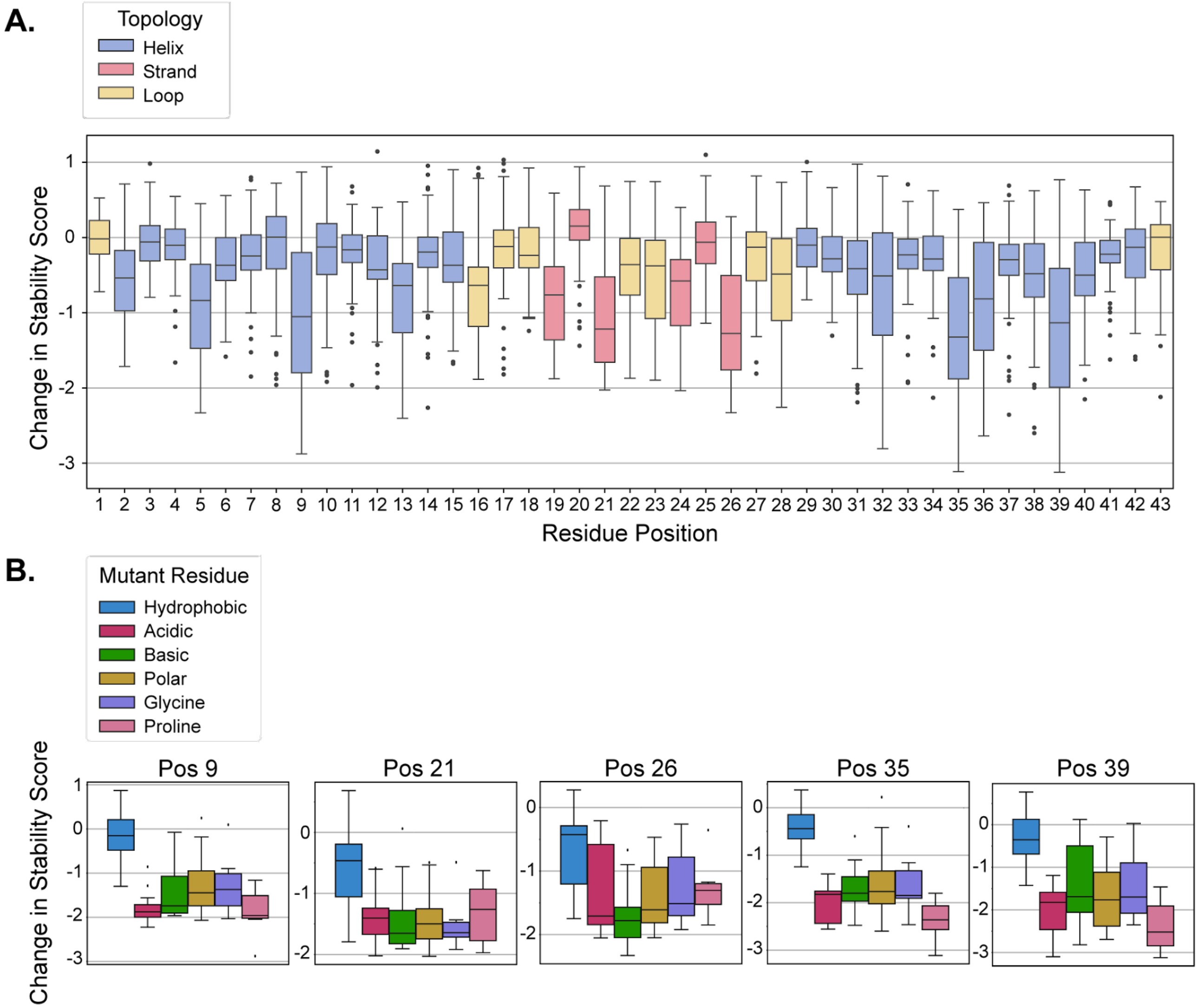


**Fig. S10. Impact of mutant residue type and the position of the mutant on stability**. To analyze whether certain mutations at specific positions on the αββα miniprotein are more likely to destabilize the structure, we (A) showed the distributions of the change in stability score (mutant stability - wild type stability) at each position of the six miniproteins of which we have deep mutational scanning data (Fig. S9). The top five destabilizing positions (9, 21, 26, 35, and 39) have an average change in stability score < 1 and are located in the buried core. For these positions (B) the change in stability score is broken down and shown by the type of mutant residue.

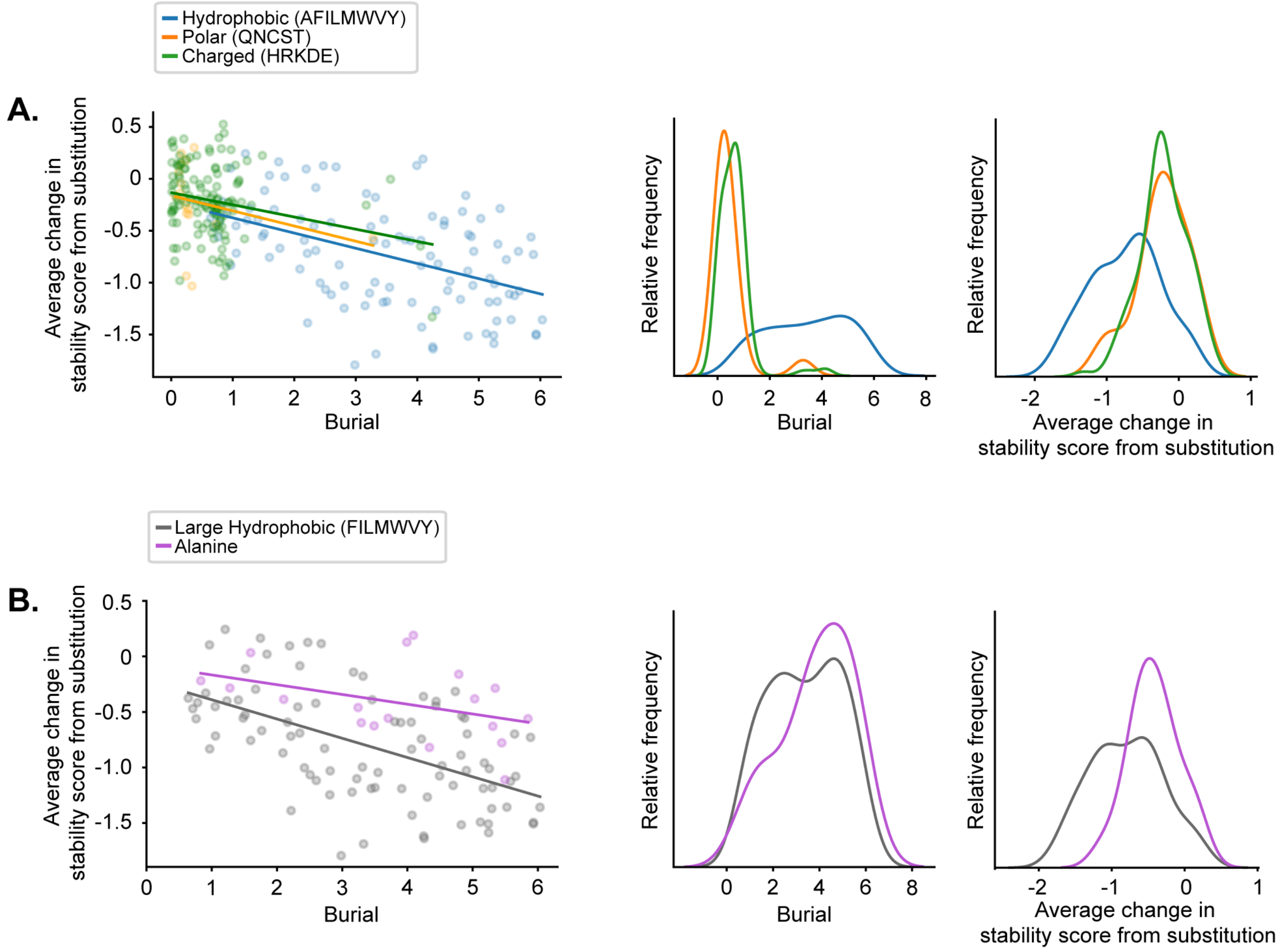


**Fig S11. Sensitivity to mutation among buried and exposed residues.** To evaluate the distribution of mutational sensitivity among buried and solvent-exposed residues, we computed the extent to which a residue in the wildtype miniprotein (that was tested in the deep mutational scanning, Fig. S9) is buried in the miniprotein (higher values indicate buried in the core, lower values indicate exposed to solvent). We compared each residue's burial value to its mutational sensitivity (average change in stability score from substitution). More negative values of the average change in stability indicate greater destabilization of the miniprotein. (A) Hydrophobic, polar, and charged residues are colored in blue, orange, and green, respectively; (B) Large hydrophobic and Alanine (small) are colored in gray and purple, respectively. The burial for a given residue was computed from the backbone coordinates of the designed structure by summing the number of CA atoms in a cone projecting out from the residue's CA-CB vector. The script is provided in the supplementary information.

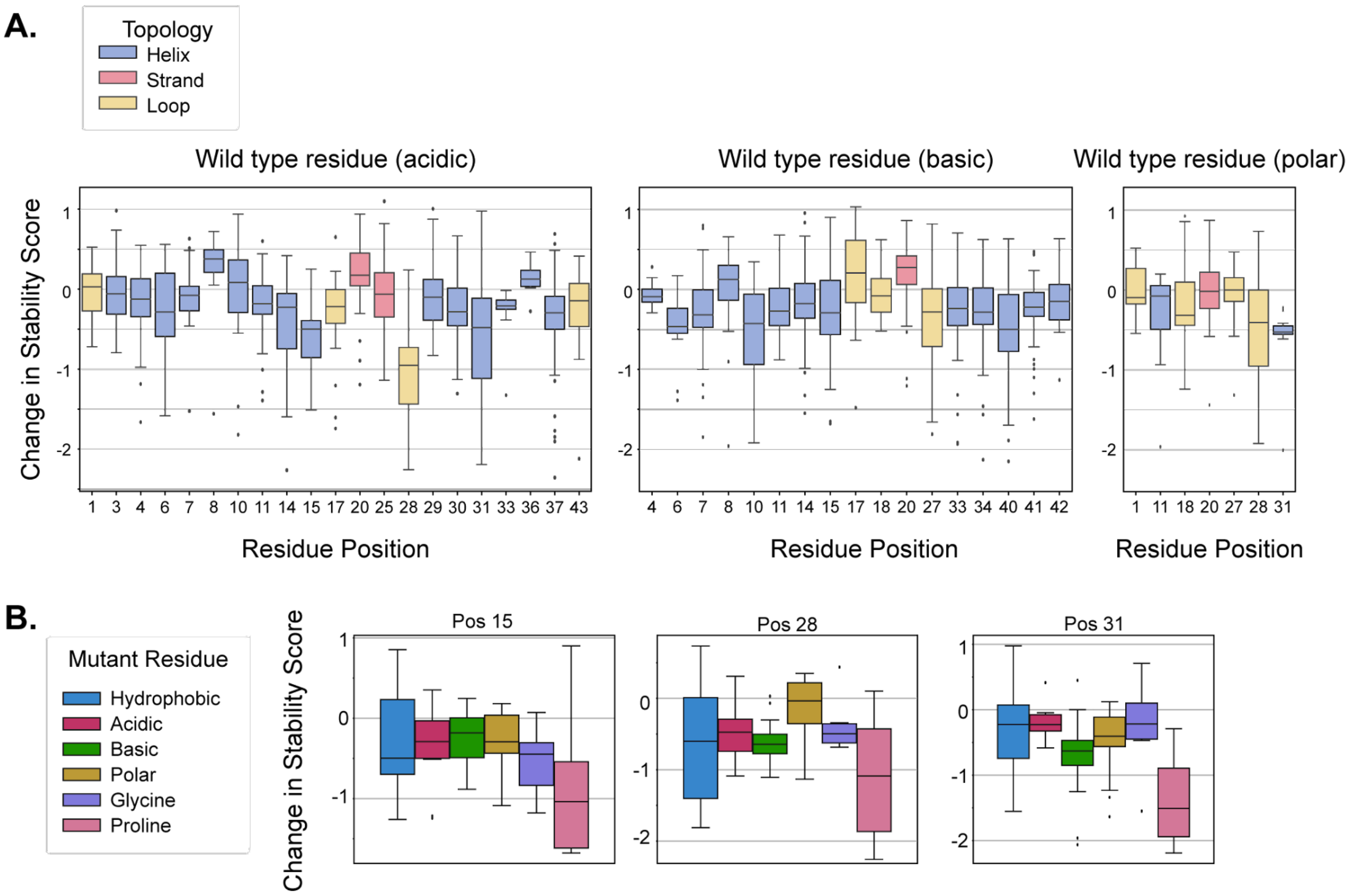


**Fig. S12. Impact of mutant residue type and the position of the mutant on stability**. To analyze whether certain mutations at specific positions on the backbone are more likely to destabilize the miniprotein, we (A) showed the distributions of the change in stability score (mutant stability - wild type stability) at each position of the six miniproteins of which we have deep mutational scanning data. The top five destabilizing positions (9, 21, 26, 35, and 36) had an average change in stability score < 1. (B) For positions 9, 21, 26, 35, and 39, the change in stability score are shown by the type of mutant residue.

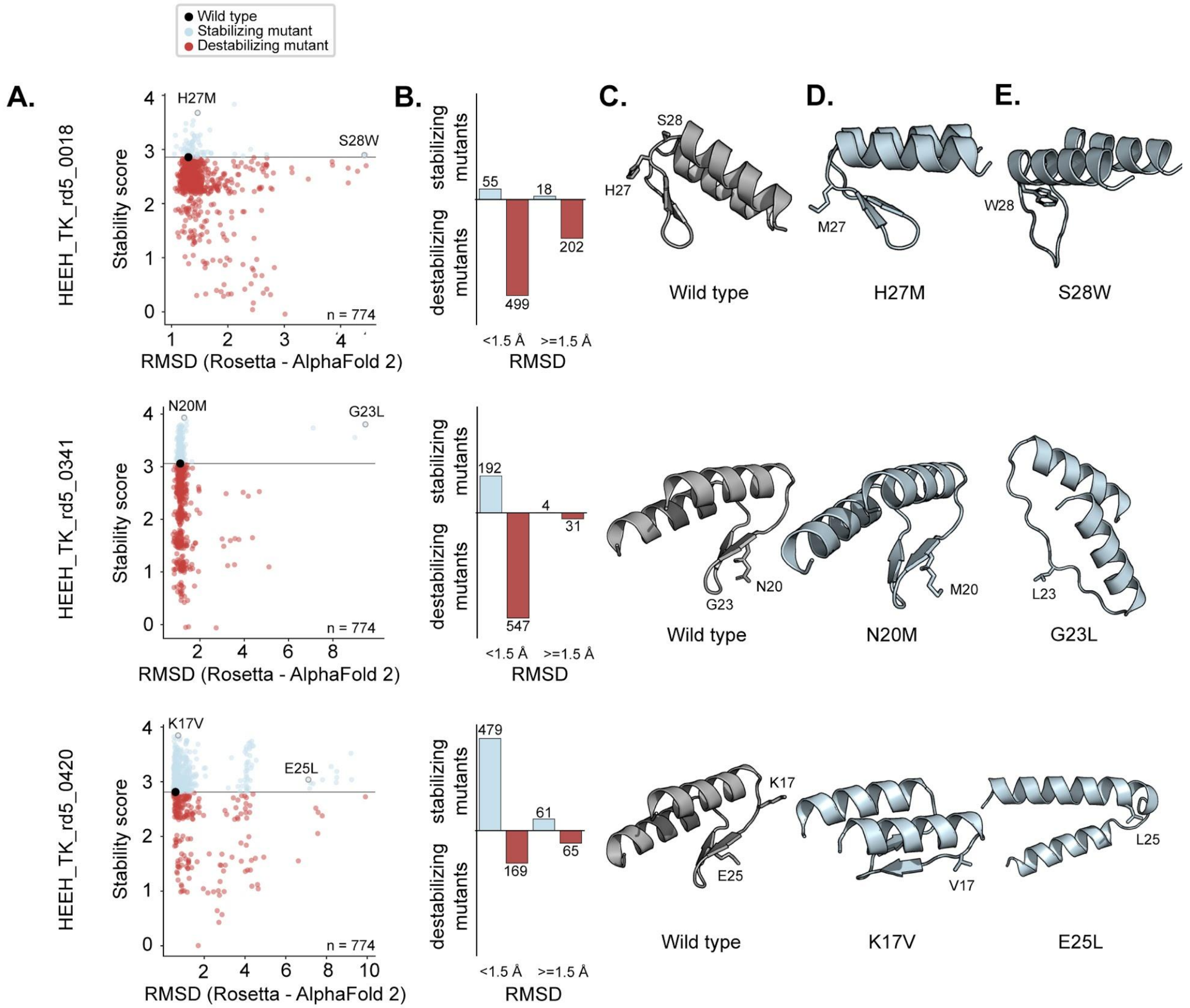

Wild type
Stabilizing mutant
Destabilizing mutant
A.
B.
C.
D.
E.
HEEH_TK_rd5_0018
Stability score
RMSD (Rosetta - AlphaFold 2)
H27M
S28W
n = 774
stabilizing mutants
destabilizing mutants
55
18
499
202
<1.5 Å
>=1.5 Å
RMSD
S28
H27
M27
W28
Wild type
H27M
S28W
HEEH_TK_rd5_0341
N20M
G23L
192
4
547
31
G23
N20
M20
L23
N20M
G23L
HEEH_TK_rd5_0420
K17V
E25L
479
61
169
65
K17
E25
V17
L25
K17V
E25L

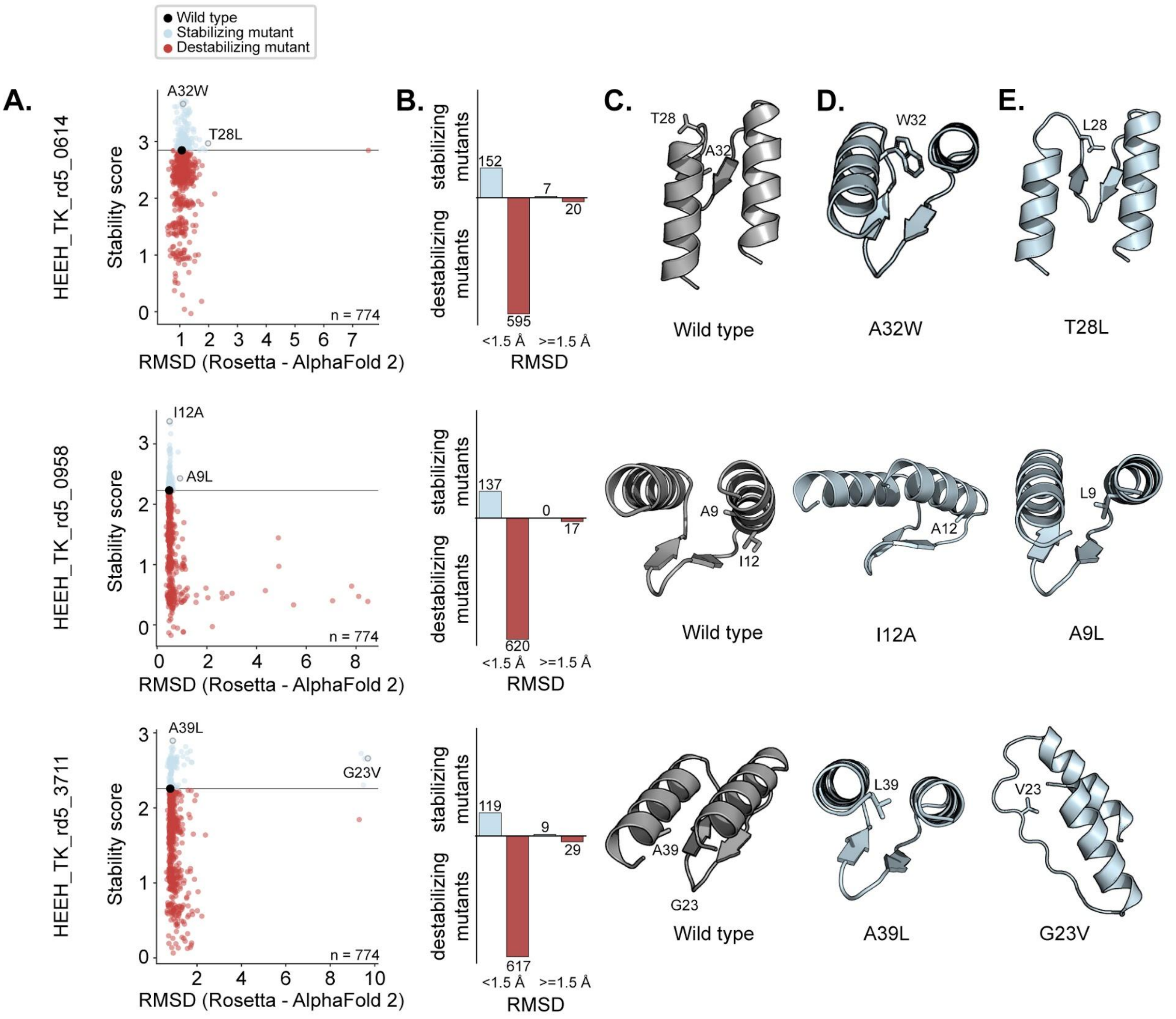


**Fig. S13. Agreement between wildtype Rosetta design models and mutant AlphaFold 2-predicted structures.** We compared how well the structure of a wildtype design (Rosetta design model) agrees with all possible mutants (whose structures are predicted by AlphaFold 2). (A) The stability score of all mutants and its corresponding wildtype are plotted by their structural agreement (RMSD); stabilizing mutants are colored in blue whereas destabilizing mutants are colored in red. (B) Stabilizing and destabilizing mutants separated by RMSD < 1.5 Å or > 1.5 Å. Cartoons of a (C) wildtype design (Rosetta design model), (D) a stabilizing mutant (AlphaFold 2-predicted structure) with close structural agreement to its corresponding wildtype, and (E) a stabilizing mutant (AlphaFold 2-predicted structure) with structural agreement that is less than (C).

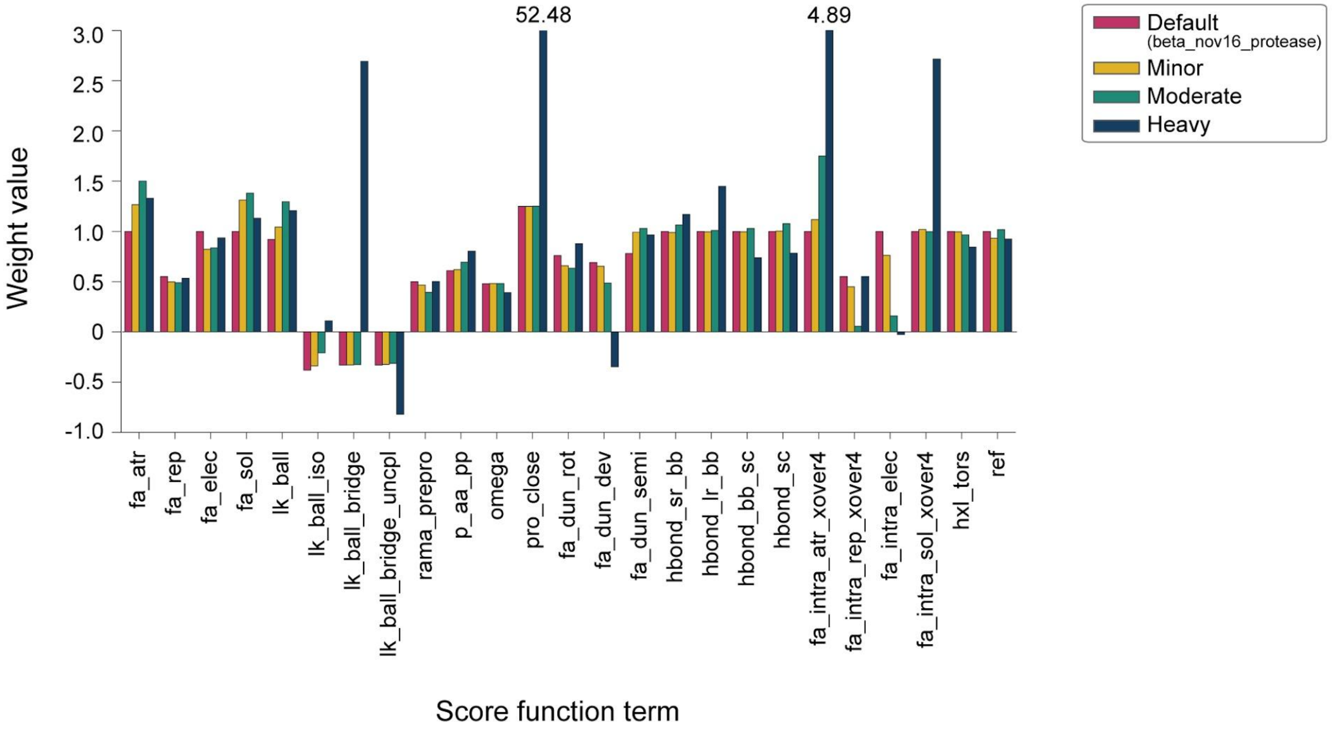


**Fig. S14. Re-weighted values of Rosetta energy score terms.** We used three differently weighted values (Minor, Moderate, and Heavy) in addition to the default values of the Rosetta score terms when designing diversity-oriented αββα miniproteins. The values of each score term are presented as a barplot. In the reweighting ridge regression, the Minor set used regularization strength alpha=200,000, Moderate used alpha=20,000, and "Heavy" used alpha=0.1.

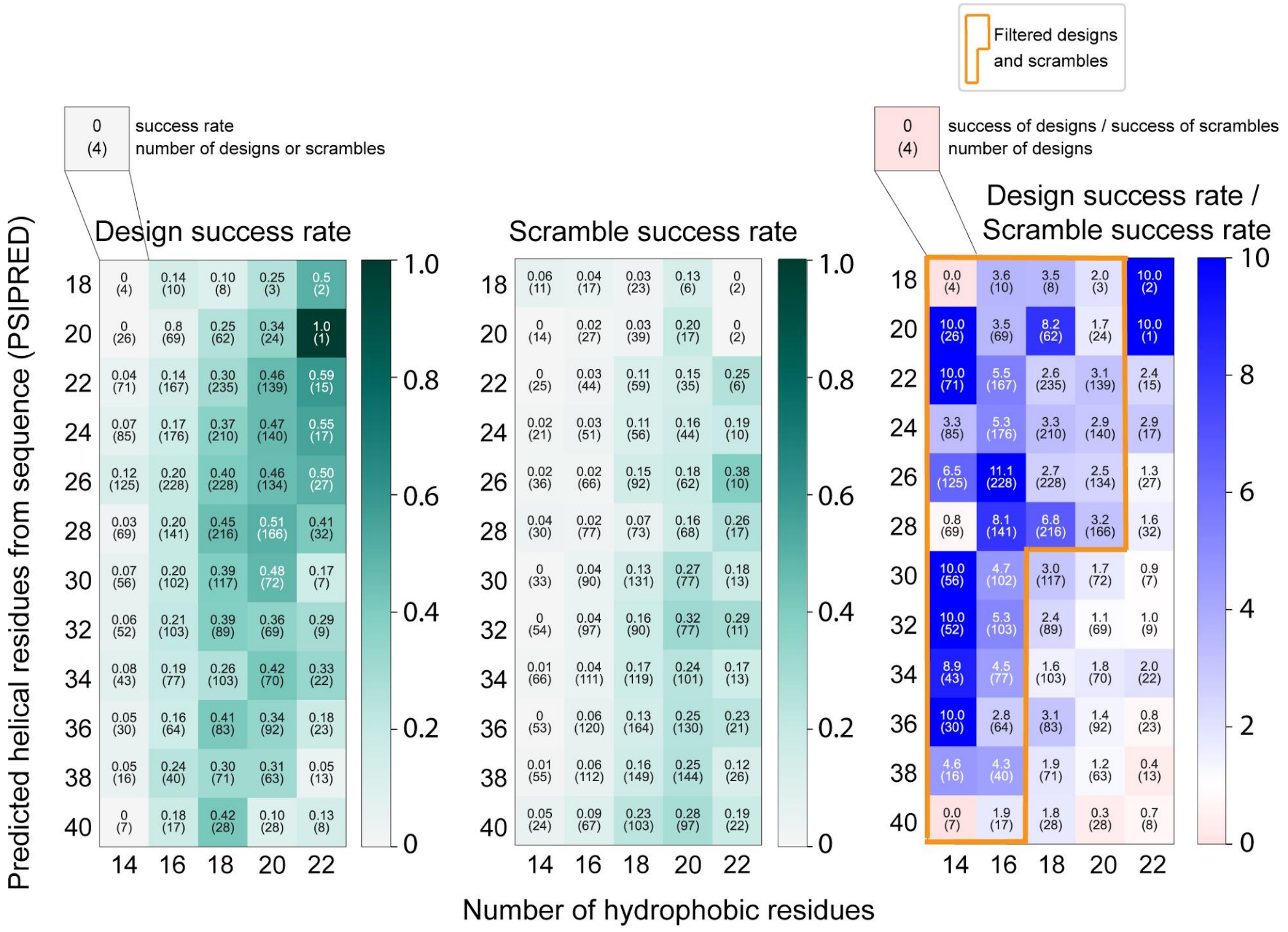


**Fig. S15. Filtering strategy for diversity-oriented designs.** In order to rule out sequences that might be "false positives" (i.e. sequences that are stable even without a specific designed structure), we examined the success rates of designs (left) and scrambled sequences (middle) after dividing them into bins according to their number of hydrophobic residues and number of residues predicted to be helical by PSIPRED (52). We minimized false positives by filtering out regions where the scrambled success rate was high (outside the orange box).

A.

| Loop 1 | Loop 2 | Loop 3 | |
|---|---|---|---|
| AABA | AAAG | AA | BBAB |
| **AGBB** | AAG | AAB | BBAE |
| AGGA | AGAB | AAGB | BBB |
| BAA | BG | **AB** | BBBB |
| BOB | BGA | ABA | BBBE |
| G | BGG | ABAA | BBE |
| GAB | BGGB | ABAB | BBEA |
| GBBA | **EA** | ABB | BBEB |
| GBBB | GA | ABBB | BBGA |
| GBBE | GEA | ABBE | **BBGB** |
| GBBG | GEB | AE | BEAA |
| GGBB | **GG** | AEA | BEB |
| BAAB | GGA | AEBB | BGB |
| BABB | GGB | AGB | E |
| GEB | GGG | AGBA | EAAB |
| AAGB | OAB | B | EAB |
| ABAB | AA | BAAA | EBGB |
| AGB | BAAB | **BAAB** | EEB |
| AGBA | BABB | BAAG | GAGB |
| BAB | BGB | BAB | GB |
| BBB | | BABA | GBA |
| GB | | BABB | GBAB |
| GBA | | BABG | GBB |
| GBAB | | BAE | |
| **GBB** | | BB | |

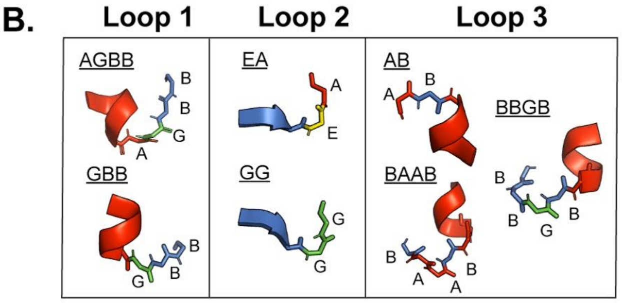


**Fig. S16. Loop patterning in diversity-oriented designs.** (A) A list of all unique loop structures (written in ABEGO notation) observed in the diversity-oriented designs. Loop structures in bold indicate that > 300 designs with that particular structure were identified in our dataset. As a result, these loop structures were selected as inputs to a topology-focused regression model (see Fig. 4F-G). (B) Cartoons visualizing the seven most common loop patterns in our data.

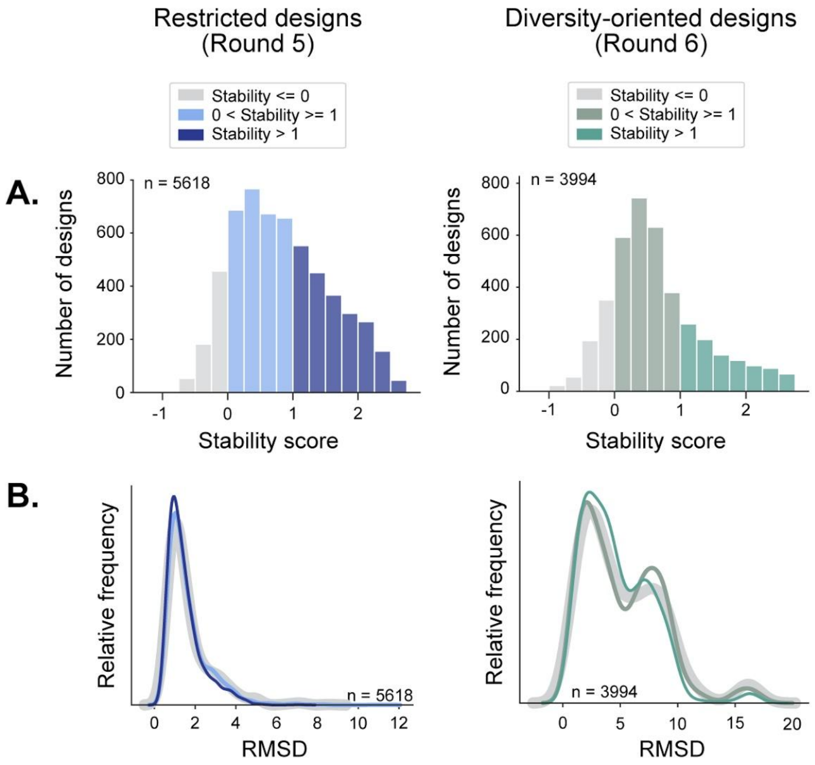


**Fig. S17. Structure agreement between AlphaFold 2 and Rosetta design models of varying stability.** (A) The distribution of αββα miniprotein stability scores, divided into three overall stability levels. (B) For each stability level, we show the distribution of structural agreement (RMSD) between Rosetta designs models and AlphaFold 2-predicted structures. Each level of stability has a similar distribution of RMSDs.

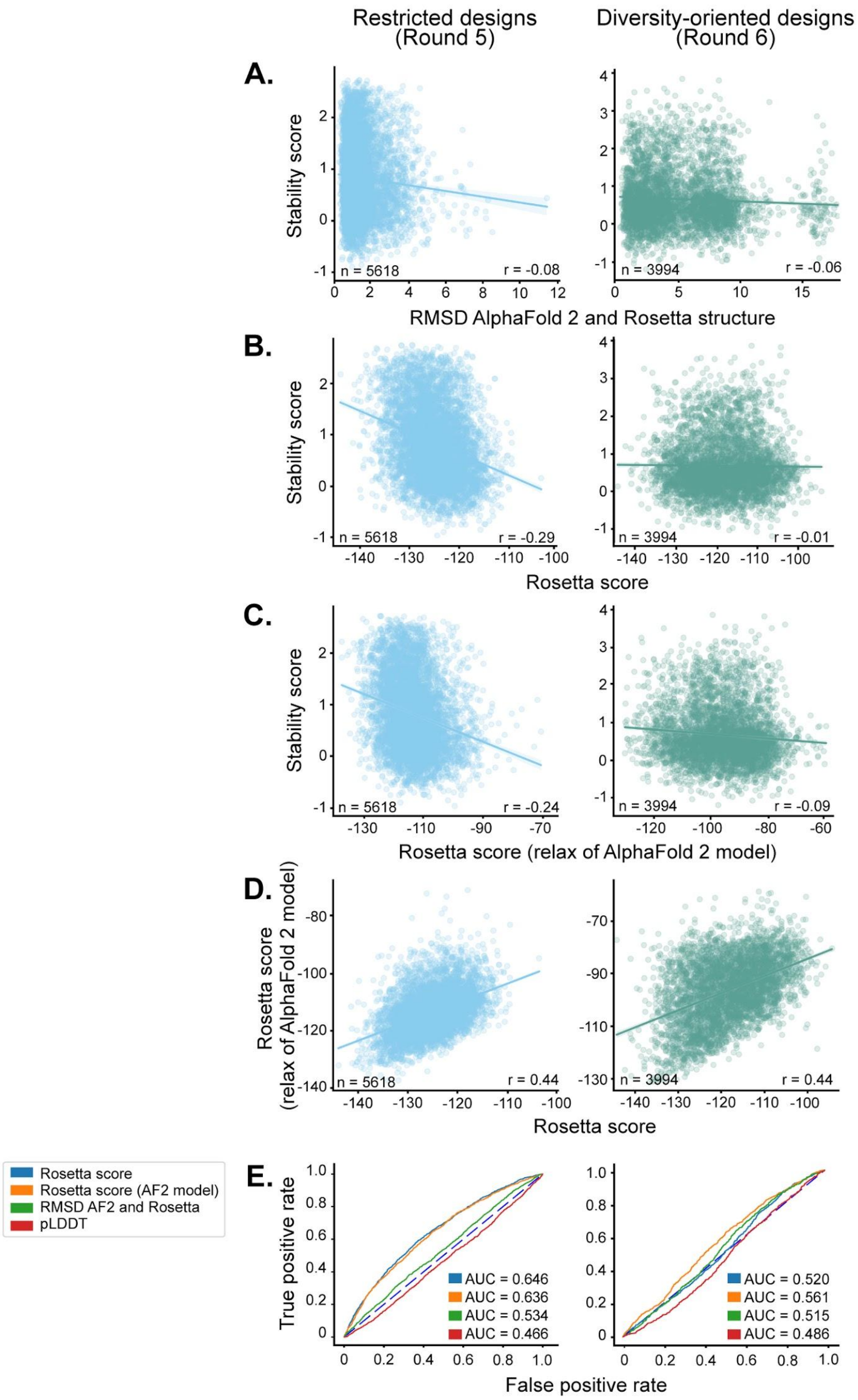

Restricted designs (Round 5)
Diversity-oriented designs (Round 6)
A.
Stability score
n = 5618
r = -0.08
n = 3994
r = -0.06
RMSD AlphaFold 2 and Rosetta structure
B.
Stability score
n = 5618
r = -0.29
n = 3994
r = -0.01
Rosetta score
C.
Stability score
n = 5618
r = -0.24
n = 3994
r = -0.09
Rosetta score (relax of AlphaFold 2 model)
D.
Rosetta score (relax of AlphaFold 2 model)
n = 5618
r = 0.44
n = 3994
r = 0.44
Rosetta score
E.
Rosetta score
Rosetta score (AF2 model)
RMSD AF2 and Rosetta
pLDDT
True positive rate
AUC = 0.646
AUC = 0.636
AUC = 0.534
AUC = 0.466
AUC = 0.520
AUC = 0.561
AUC = 0.515
AUC = 0.486
False positive rate

**Fig. S18. Comparison of Rosetta design models and AlphaFold 2-predicted structures in predicting stability.** Structures of all αββα miniproteins were predicted by AlphaFold 2, and the predicted model with the lowest RMSD to the designed structure was used for further analysis. The experimental stability scores are compared to the (A) structural agreement (RMSD) between AlphaFold 2-predicted models and Rosetta design models, (B) Rosetta energy determined from the Rosetta design models, (C) and Rosetta energy determined from the AlphaFold 2-predicted structures. (D) Rosetta energy scores determined from the Rosetta design models and AlphaFold 2-predicted models are compared to each other. (E) ROC curves to classify αββα miniproteins as stable (stability score $\geq 1$) or unstable based on scores determined from the Rosetta model (blue), AlphaFold 2-predicted structure (orange), the agreement between the Rosetta model and AlphaFold 2 structure (green), and the AlphaFold 2 confidence score (red).

### Supplementary Tables

**Table S1. Selected αββα designs for circular dichroism, thermal denaturation, and deep mutational scanning.**

| Design | Sequence (Helix, Strand) | Hydrophobicity |
|---|---|---|
| HEEH_TK_rd5_0958 | DIEEIEKKARKILEKGDSIEIAGFEVRDEEDLKKILEWLRRHG | 347 |
| HEEH_TK_rd5_3711 | SWEDLERLAREALERGETIHILGFEIRSEEDAKKFAEWARRWE | 842 |
| HEEH_TK_rd5_0341 | DLEELEEDLKQALREGRKVNILGIEVTTEEQARRLIEFLRRFI | 968 |
| HEEH_TK_rd5_0614 | DLEKLRELLEDALRKGITIRFAGIEVKTEEEAERLLEWLKRKL | 1018 |
| HEEH_TK_rd5_0018 | SFEELIKLIEDLLRKGDHINILGFEVHSEEEARRLIEWLRRAA | 1174 |
| HEEH_TK_rd5_0420 | SLEELLKLAEEALKRGKTIRILGFEISSEEALRRFEEWLRRFI | 1258 |

* gray and blue boxes indicate residues that are in the helices and β-strands, respectively; hydrophobic residues are colored in orange. Hydrophobicity values quantified based on (6).

**Table S2. NMR restraints, structural statistics, quality scores, rotational correlation times and hydrodynamic radii for HEEH_TK_rd5_0341 and HEEH_TK_rd5_0958**

| **Design ID**[a] | HEEH_TK_rd5_0341 | HEEH_TK_rd5_0958 |
|---|---|---|
| **PDBID** | 7T2F | 8DOA |
| **BMRBID** | 30974 | 31033 |
| **Relaxation-derived Dynamics** | | |
| Rotational correlation time, τc (ns) | 11.9 | 6.1 |
| Hydrodynamic radius, rH (nm) | 2.15 | 1.72 |
| **NMR restraints:** | | |
| Total NOEs | 1564 | 546 |
| Intra-residual | 437 | 198 |
| Sequential (i – j = 1) | 465 | 151 |
| Medium-range (1 < i – j < 5) | 333 | 98 |
| Long-range (i – j ≥ 5) | 275 | 99 |
| Inter-molecular | 54 | N/A |
| Hydrogen Bonds | 44 | N/A |
| Dihedral Angles: | | |
| φ | 78 | 42 |
| ψ | 78 | 42 |
| **Structural Statistics:** | | |
| r.m.s.d. from experimental restraints: | | |
| Distance restraints (Å) | 0.0197 ± 0.002 | 0.0072 ± 0.0016 |
| Dihedral angle restraints (°) | 0.280 ± 0.14 | 0.463 ± 0.19 |
| Violations in the NMR ensemblea: | | |
| Max. distance restraint violation (Å) | 0.503 | < 0.3 |
| Max. dihedral restraint violation (°) | 6.2 | 5.2 |
| r.m.s.d. from idealized geometry: | | |
| Bond lengths (Å) | 0.0147 ± 0.0002 | 0.0140 ± 0.0004 |
| Bond angles (°) | 0.99 ± 0.017 | 0.90 ± 0.027 |
| Impropers (°) | 1.65 ± 0.09 | 1.59 ± 0.15 |
| Average pair-wise r.m.s.d. (Å)[b]: | | |
| Heavy | 1.2 | 1.5 |
| Backbone | 0.6 | 0.7 |
| **Structure quality scores:** | | |
| Ramachandran plot (%)[b,c] | | |
| Most favored | 92.7 | 92.7 |
| Additionally allowed | 7.3 | 7.3 |
| Generously allowed | 0.0 | 0.0 |
| Disallowed | 0.0 | 0.0 |
| Structural Quality Factors (raw/Z-scores)[d] | | |
| Procheck (phi/psi) | 0.00/0.31 | 0.13/0.83 |
| Procheck (all) | -0.18/-1.06 | -0.04/-0.24 |
| Molprobity clash | 15.72/-1.17 | 8.71/0.03 |

[a]The NMR ensemble consists of the 20 lowest energy structures out of 100 calculated; [b]Calculated for residues 22 to 62 inclusive for HEEH_TK_rd5_0958, and residues 23-36, 40-42, 45-47 and 50-62 inclusive, for HEEH_TK_rd5_0341; [c]Based on Procheck analysis (7); [d]Calculated using the PSVS server (8) (https://montelionelab.chem.rpi.edu/PSVS/).

**Table S3. Coefficients of a ten-feature linear regression model.**

| Feature | Coefficient Δ Stability Score (per 1 σ) | Coefficient Δ Stability Score (per contact) |
|---|---|---|
| Large nonpolar count (FILMWY) | 0.256 | 0.161 |
| Nonpolar residue-residue contacts | 0.189 | 0.048 |
| Local sequence-structure propensity | 0.129 | 0.129 |
| Ser or Thr in helix caps | 0.086 | 0.125 |
| Glu-Arg residue-residue contacts | 0.082 | 0.038 |
| Nonpolar residue at design ends | 0.070 | 0.078 |
| Favorable net charge at helix ends | 0.042 | 0.020 |
| Glu-Glu residue-residue contacts | -0.023 | 0.086 |
| Buried unsatisfied polar atoms | -0.058 | -0.134 |
| Increased net charge$^2$ | -0.116 | -0.085 |

**Table S4. Mean correlation coefficients and 95% confidence intervals of linear regression models.**

| Model | Mean r | 95% CI |
|---|---|---|
| 10 features | 0.642 | (0.628, 0.657) |
| 10 features + 25 score terms | 0.673 | (0.659, 0.687) |
| 10 features + 25 shuffled score terms | 0.646 | (0.630, 0.662) |
| 10 features + shuffled stability scores | 0.059 | (0.038, 0.081 |
| 10 features + 25 shuffled score terms with shuffled stability scores | 0.107 | (0.083, 0.130) |

**Table S5. Reweighted Rosetta score functions applied to miniprotein designs of varying topologies**

| **Spearman's Correlation** | | | | | |
|---|---|---|---|---|---|
| Rounds 1-4 | **ααα** | **αββα** | **βαββ** | **ββαββ** | **Score function** |
| | -0.505 | -0.089 | -0.390 | -0.628 | beta_nov16_protease |
| | -0.610 | -0.172 | -0.479 | -0.631 | minor |
| | -0.605 | -0.165 | -0.482 | -0.624 | moderate |
| **ROC** | | | | | |
| Rounds 1-4 | 0.767 | 0.636 | 0.710 | 0.901 | beta_nov16_protease |
| | 0.824 | 0.672 | 0.767 | 0.911 | minor |
| | 0.821 | 0.681 | 0.769 | 0.908 | moderate |
| **Spearman's Correlation** | | | | | |
| Round 4 only | -0.058 | -0.070 | -0.189 | -0.375 | beta_nov16_protease |
| | -0.185 | -0.153 | -0.262 | -0.473 | minor |
| | -0.213 | -0.147 | -0.268 | -0.431 | moderate |
| **ROC** | | | | | |
| | 0.515 | 0.602 | 0.596 | 0.714 | beta_nov16_protease |
| | 0.592 | 0.654 | 0.644 | 0.758 | minor |
| | 0.609 | 0.647 | 0.656 | 0.734 | moderate |

Re-weighted energy functions (minor and moderate) were used to calculate the predicted scores of previously published protein designs (four different topologies). The Spearman's correlation coefficient was determined based on comparing the experimental stability score vs. the predicted stability score. beta_nov16_protease is the score function used previously (1). Negative Spearman's correlations are expected because more negative energy scores and more positive stability scores both imply greater stability.
